# Rapid, concerted switching of the neural code in inferotemporal cortex

**DOI:** 10.1101/2023.12.06.570341

**Authors:** Yuelin Shi, Dasheng Bi, Janis K. Hesse, Frank F. Lanfranchi, Shi Chen, Doris Y. Tsao

## Abstract

A fundamental paradigm in neuroscience is the concept of neural coding through tuning functions^1^. According to this idea, neurons encode stimuli through fixed mappings of stimulus features to firing rates. Here, we report that the tuning of visual neurons can rapidly and coherently change across a population to attend to a whole and its parts. We set out to investigate a longstanding debate concerning whether inferotemporal (IT) cortex uses a specialized code for representing specific types of objects or whether it uses a general code that applies to any object. We found that face cells in macaque IT cortex initially adopted a general code optimized for face detection. But following a rapid, concerted population event lasting < 20 ms, the neural code transformed into a face-specific one with two striking properties: (i) response gradients to principal detection-related dimensions reversed direction, and (ii) new tuning developed to multiple higher feature space dimensions supporting fine face discrimination. These dynamics were face specific and did not occur in response to objects. Overall, these results show that, for faces, face cells shift from detection to discrimination by switching from an object-general code to a face-specific code. More broadly, our results suggest a novel mechanism for neural representation: concerted, stimulus-dependent switching of the neural code used by a cortical area.

## Introduction

A core challenge of visual neuroscience is to understand how populations of neurons encode visual stimuli. Over the past decade, major progress on this question has been made in macaque inferotemporal (IT) cortex^2–6^, a large brain region dedicated to high-level object recognition^7–9^. IT cortex is anatomically organized into a set of parallel networks, each specialized for encoding specific aspects of object shape and color^6,10–13^. One of these networks, the macaque face patch system, has served as a crucible for testing competing theories of high-level object coding^14–16^. Here, we exploit this system to resolve a central debate about the coding mechanism used by IT cortex: does it rely on specialized mechanisms for processing specific types of objects^17–19^, or does it rely on general mechanisms that process all types of objects in the same way^20–22^?

According to one theory, face processing is “domain specific,” i.e., reliant upon specialized mechanisms dedicated to faces^17,23^. This idea originated from the behavioral observation that humans are much better at discriminating face parts presented in the context of a whole face^24,25^ (**Fig. 1a**, top right inset), as well as the neuropsychological observation of “prosopagnosia,” a highly specific deficit in face recognition that can be caused by a focal lesion of the temporal lobe^26,27^. These observations have led to the suggestion that a face detection gate–a switch determining whether an image is a face or not–is implemented prior to detailed feature processing^23^ (**Fig. 1a**, top).

**Figure 1:**
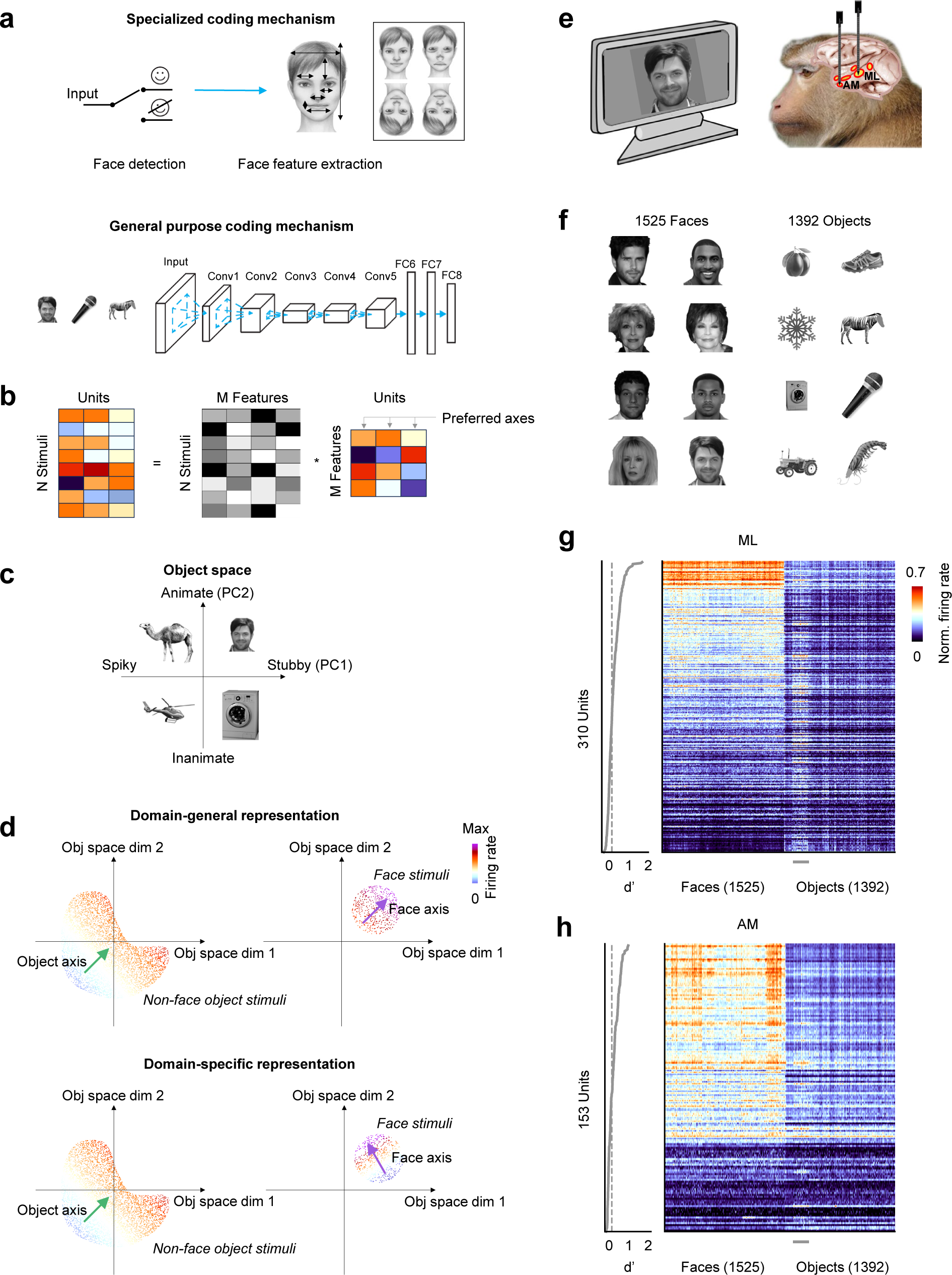
Two contrasting models for face processing. [Note: all human faces in this figure have been replaced by synthetically generated faces using a diffusion model^57^ due to biorxiv policy on displaying human faces.] **a.** Top: schematic of a specialized coding mechanism. Extraction of detailed face features is preceded by a face detection step, a gate that ensures only upright faces undergo detailed processing. Inset, top right: the Thacher illusion, illustrating that face features are much better perceived when embedded in an upright face. Bottom: schematic of a general-purpose coding mechanism. Face cells are modeled by units in a late layer of deep neural network that perform the same computation on faces and objects. **b.** Computation of preferred axes of units via linear regression of face/object responses to stimulus features derived from a DNN representation. **c.** A 2D general object space can be computed by principal components analysis (PCA) of a deep neural network representation of faces and objects^6^. Faces generally cluster in one quadrant. **d.** Schematic illustrating predictions of domain-general (top) vs. domain-specific (bottom) models of face processing. In the top left plot, each dot corresponds to the projection of one *object* image onto the first two dimensions of object space, and the color of the dot indicates the firing rate of a cell elicited by the image. The green arrow is the cell’s object axis and points in the direction of the response gradient. In the top right plot, each dot now represents projection of one *face* image onto the first two dimensions of the same object space. The purple arrow indicates the cell’s face axis. If cells use a domain-general mechanism (top), the face and object axes should align. On the other hand, if cells use a domain-specific mechanism (bottom), then the face and object axes should differ. **e.** Schematic of experiment. We recorded simultaneously from two faces patches, ML and AM, with NHP Neuropixels probes, while monkeys fixated visual stimuli. **f.** In the main experiment, a large set of human faces (left) and non-face objects (right) were presented. **g.** Right: mean responses of all visually-responsive cells in ML (N = 310) to each face and object stimulus, computed using a time window of 50-220 ms and normalized to the range [0 1] for each cell. The small gray horizontal bar at bottom indicates images of animals with grayed-out heads. Left: face selectivity d’ for each cell. The dotted vertical line marks d’ = 0.2, which we used as our threshold for identifying face-selective units. **h.** Same as (g), for AM cells (N = 153).

Multiple findings from the macaque face patch system support the idea that face processing is domain specific. First, the very existence of a system of patches in which a majority of cells respond more strongly to faces than to other objects suggests a specialized mechanism for processing faces^15,22,28,29^. Second, electrical stimulation of face patches strongly distorts monkeys’ perception of facial identity, while having little effect on perception of clearly non-face objects^30^. Third, cells in face patches are exquisitely sensitive to the geometry, contrast, and texture of faces^3,14,31^. All these findings suggest a special role for the macaque face patch system in representing faces.

However, a competing theory argues that the computation performed by face patches is domain general and not specific to faces^20,22^. Several studies have found that general-purpose deep neural networks (DNNs) trained on object categorization provide an effective model for cells across all of IT cortex, including within face patches^2,6,22^ (**Fig. 1a**, bottom). Specifically, IT cells can be modeled as computing linear combinations of features represented in late layers of these networks: 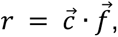, where *r* is the response of the IT cell, 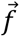 is a vector of object features given by the DNN representation, and 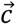 defines the “preferred axis” of the cell^2,6,22^ (**Fig. 1b**). According to this picture, the DNN feature representation defines a general “object space” in which all objects, including faces, live (**Fig. 1c**). Intuitively, the preferred axis is the direction of the response gradient of a cell (i.e., the vector pointing from stimuli in the object space that elicit weak responses to stimuli that elicit strong responses). In this framework, face cells–just like other IT cells–encode objects by projecting incoming stimuli onto their preferred axis agnostic to their category^6,22^.

Consistent with the idea that IT computations are domain general, it has been found that IT cortex contains a topographic map of the object space defined by the top two principal components of the DNN feature representation^6,32^ (**Fig. 1c**). Faces are clustered in one quadrant of this space, and preferred axes of face cells consistently point towards this quadrant, while preferred axes of cells in other IT patches point in other directions^6^. Thus one interpretation is that face cells are not special: just like other IT cells, they are performing domain-general axis projection. This domain-general theory of face patch coding predicts that each cell should have a single preferred axis, i.e., a single response gradient spanning the entire general object space. If true, this would imply that one can use either faces or objects to map this gradient, and both methods should yield the same preferred axis^6,22^ (**Fig. 1d**, top).

In contrast, the domain-specific theory of face patch coding argues that the processes underlying fine facial feature extraction are specialized for faces and are triggered only following successful detection of an upright face (**Fig. 1a**, top). Accordingly, this theory predicts that the preferred axis of a cell mapped using faces is specially tuned for face representation and is likely different from that mapped using objects (**Fig. 1d**, bottom).

To investigate these competing theories and clarify the representational logic of face cells, here we recorded responses of face cells in macaque face patches to a large set of faces and objects. We parameterized the stimuli using a deep neural network, AlexNet layer fc6^6,33,34^, creating a general object space capturing face and object variations. Prior work has shown that AlexNet provides an effective model for IT cells generally^6,35^ as well as face cells specifically^34^, and its performance is representative of a large class of feedforward DNNs^6,36^. Comparison of the encoding axes for faces vs. those for objects unexpectedly revealed a new form of neural computation: *dynamic, stimulus-dependent switching of the neural code used to represent a visual stimulus.* Results of multiple analyses converged on this conclusion. First, the preferred axes of face cells in the general object space computed using responses to objects were uncorrelated with those computed using responses to faces; the former consistently pointed in the face quadrant of the general object space, supporting robust face detection, while the latter pointed in seemingly diverse directions. Second, analysis of axis dynamics using a time-varying window revealed that the face encoding axis in the general object space was actually *initially aligned* to the object encoding axis, resembling domain general processing mechanisms. But then, a drastic switch occurred in the neural code for faces at ∼100 ms. At this latency, the face encoding axis in low dimensions of the general object space reversed direction and responses sparsened; simultaneously, robust new tuning developed for multiple face features. Strikingly, these dynamics were face specific and did not occur in response to objects, demonstrating domain-specific processing mechanisms. The new tuning to facial features enabled improved reconstructions of face identity from neural activity. *Overall, these results show that face cells achieve domain specificity over time through a rapid, concerted change in the neural code they use to represent a face,* resolving a longstanding debate on domain specific vs. domain general processing^17–22^. Our findings challenge the notion that object recognition can be explained by largely feedforward processes^37^, instead revealing an essential role for recurrent dynamics in core visual processing. More broadly, the results reveal a novel mechanism for neural representation: concerted, stimulus-driven switching of the neural code used by a cortical area.

## Results

We identified face patches ML (middle lateral) and AM (anterior medial) in two animals using fMRI^24^ (see Methods). We then targeted NHP Neuropixels probes^38^ to ML in two monkeys and AM in one monkey (**Fig. 1e**, **Extended Data Fig. 1**) and recorded responses to 1525 grayscale images of real human faces and 1392 grayscale images of diverse non-face objects (**Fig. 1f**). Each stimulus was presented for 150 ms ON and 150 ms OFF while animals passively fixated. In the main figures below, we show results from face patch ML of monkey A (results were qualitatively similar for ML in monkey J, **Extended Data Fig. 2**, and AM in monkey A, **Extended Data Figs. 6**, **9**, **11**).

### Face cells use distinct axes to represent faces and objects

Cells in both ML and AM responded more to faces compared to objects (**Fig. 1g, h**; **Extended Data Fig. 3**), with 54.5% of ML cells (N = 310) and 51.6% of AM cells (N = 153) having a face selectivity d’ ≥ 0.2 (see Methods). These percentages are lower than what has been reported using single tungsten recordings^15,28^; since Neuropixels probes span a depth of 3.84 mm, part of the probe was outside the face patch.

We first passed a combined set of face and object stimuli through AlexNet and performed principal components analysis (PCA) on the fc6 layer embedding to extract a general 60-d feature space capturing features of both faces and objects^6^. This general object space could explain 80.9% (61.6%) of the variance in the fc6 features of the object (face) stimuli. For each face cell, we used responses to the non-face object stimuli (averaged over the time window 50-220 ms) to compute a preferred axis, the *object axis*, by linearly regressing responses of cells to the 60-d feature vectors corresponding to different objects (**Fig. 1b**). Similarly, we used responses to the face stimuli to compute a preferred *face axis*. We confirmed that the object and face axes could explain significantly more variance in responses to objects and faces, respectively, compared to a shuffle control (**Extended Data Fig. 4**).

Do single face cells encode the entire object space using a single axis or do they use multiple axes (**Fig. 1d**)? As explained above, a domain-general account of face cell computation predicts that cells use a single axis^22^, and hence the face and object axes for each cell should be identical. We first compared face and object axes by visualizing them in a 2D space corresponding to the first two PCs of the 60-d feature space. In agreement with a previous study^6^, we found that the object axes of face cells consistently pointed in the upper right quadrant of the PC1-PC2 space (**Fig. 2a**, left; standard deviation of angle = 37.4°), where faces reside (**Extended Data Fig. 5a**). In contrast, the face axes of the same cells pointed in diverse directions (**Fig. 2a**, right; standard deviation of angle = 81.9°). We confirmed this using an F-test (F(1,150) = 214.7, p < 4.9*10^-37^). To ensure that the consistency of the object axes was not due to any inherent non-uniform distribution of the object stimuli (**Extended Data Fig. 5a**), we selected a subset of object stimuli that were Gaussian distributed (**Extended Data Fig. 5b**). This did not change the qualitative difference in angular distributions for object compared to face axes (**Extended Data Fig. 5c, d**), which was confirmed to be destroyed by stimulus shuffling (**Extended Data Fig. 5e, f**). A scatter plot of face and object axis weights for PC1 confirmed lack of clear correlation (**Fig. 2b**, left; r = 0.16, p = 0.04), and similarly for PC2 (**Fig. 2b**, right; r = −0.07, p=0.37). Further confirming the lack of correlation between face and object axes, the face axis could not explain any variance in the object responses, and vice versa (**Fig. 2c**).

**Figure 2:**
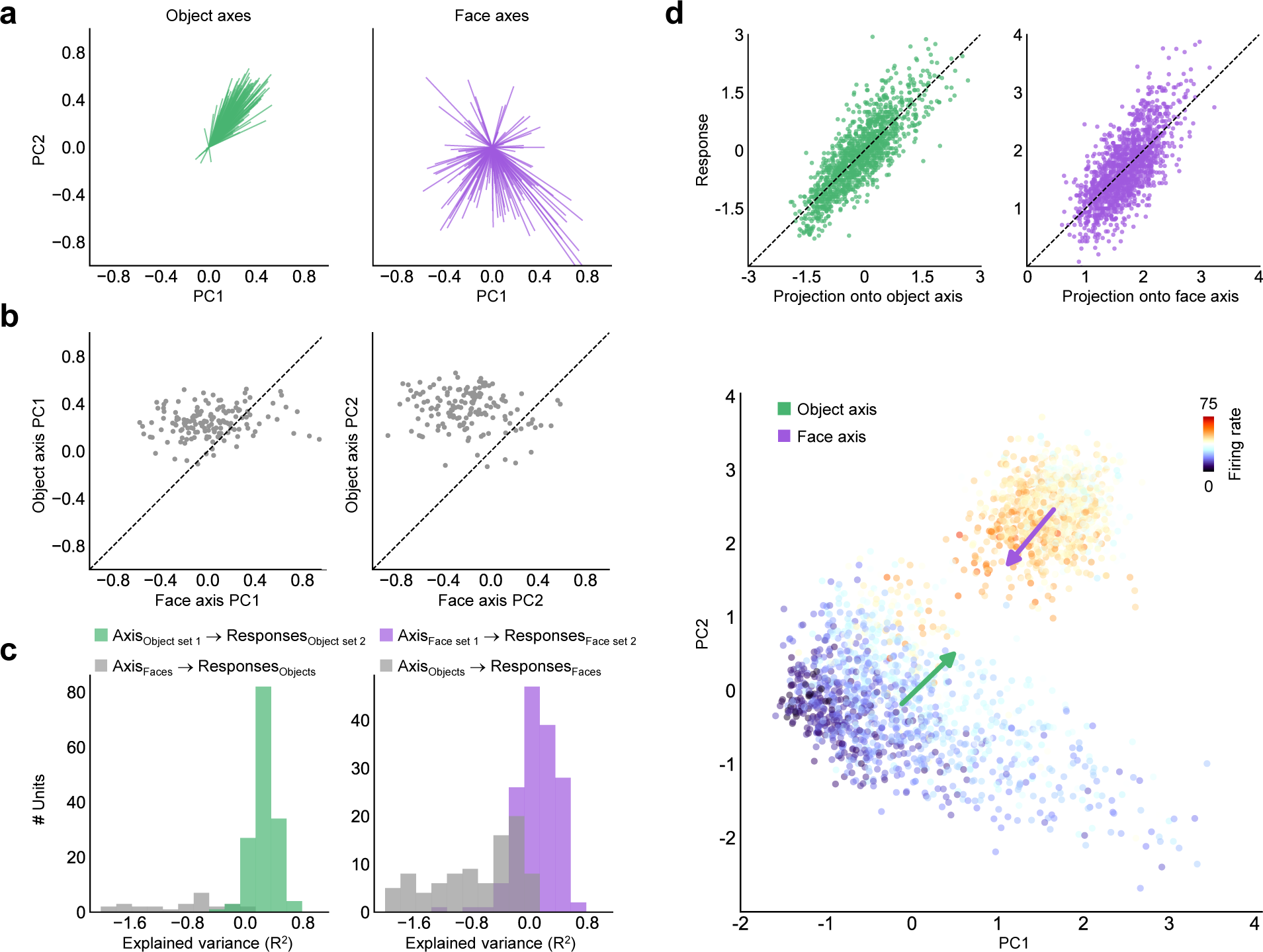
Face cells use distinct axes to represent faces and objects. **a.** Distribution of object (left, green) and face (right, purple) axes, projected onto the top two dimensions of the 60-d object space (note that faces occupy the upper right quadrant of this space, cf. Extended Data Fig. 5a) (N = 151 cells, only cells with R^2^ > 0 on test set for object and face axes are included, see Methods). **b.** Scatter plots of object vs. face axis weights for PC1 (left) and PC2 (right). **c.** Cross prediction accuracies. Left: distribution of explained variances when using (i) face axes to explain object responses (gray) or (ii) object axes computed using responses to a random object subset to explain responses to the left-out object subset (green). Right: same as left, but using the object axes to explain face responses. Units with R^2^ < −2 are not shown. **d.** Responses of an example cell with opposite tuning to objects and faces. Top Left: scatter plot showing actual responses to objects vs. predicted responses computed by projecting images onto the object axis (R^2^ = 0.62). Top right: scatter plot showing actual responses to faces vs. predicted responses computed by projecting images onto the face axis (R^2^ = 0.54). Bottom: responses of this cell to objects (large cluster in lower left) and faces (small cluster in upper right), color coded by response magnitude (cf. Extended Data Fig. 5a). The object (face) axis of the cell is indicated by the green (purple) arrow.

The lack of correlation between face and object axes was evident in the raw responses of single cells. **Fig. 2d** shows an example cell with diametrically opposite face and object axes. Responses of this cell to objects were well captured by the object axis (**Fig. 2d**, top left); similarly, responses to faces were well captured by the face axis (**Fig. 2d**, top right). Yet the two axes clearly pointed in opposite directions in the PC1-PC2 space (**Fig. 2d**, bottom). Thus it is clear from the raw responses that a single axis cannot explain the selectivity of this cell for faces and objects (cf. **Fig. 1d**).

One might wonder if the observed difference between the face and object axes depended on prior expectations. To obtain the data in **Fig. 2**, face and object stimuli were presented in separate blocks of only faces or only objects (see Methods), so prior expectations could conceivably have played a role in generating axis differences. However, we observed the same pattern of results when the face and object stimuli were randomly interleaved (**Extended Data Fig. 2a-d**). This suggests that the axis difference is not related to prior expectations/adaptation and is instead purely driven by the current stimulus.

Cells in face patch AM showed a very similar pattern of responses (**Extended Data Fig. 6**), with object axes consistently pointing towards the face quadrant and face axes pointing in diverse directions.

Could the lack of generalizability of the object axis for explaining responses to faces and vice versa (**Fig. 2c**) be due to the well-known phenomenon of increased error for out-of-distribution (OOD) generalization^22^? How would face-selective units that unambiguously use a single axis fare under the same analyses? Would the OOD problem be so severe that mapping such units, where we know the ground truth, with faces and objects would falsely reveal uncorrelated axes? To address this, we mapped responses of artificial face-selective units in AlexNet layer fc6 to the same face and object stimuli shown to the monkey. AlexNet fc6 units should have a single axis in the 60-d feature space since the feature space was computed by PCA on these units. When the analyses in **Fig. 2a-d** were applied to AlexNet units, face and object axes were indeed correlated (**Extended Data Fig. 7**), matching the ground truth and clearly differing from macaque face patch neurons.

Overall, these results suggest that single face cells encode faces and objects using distinct axes, unlike units in a DNN.

### The face axis initially aligns to the object axis before suddenly reversing

If face and object axes are distinct, where and how is the decision made regarding which axis a face cell should use? Is the choice apparent in the first spikes, or does it evolve dynamically within a face patch? Single face cells showed rich response dynamics that differed between faces and objects. For example, the cell in **Fig. 3a** gave a strong transient response to faces followed by a sustained period of firing; in response to objects, this cell showed a sustained response throughout.

**Figure 3:**
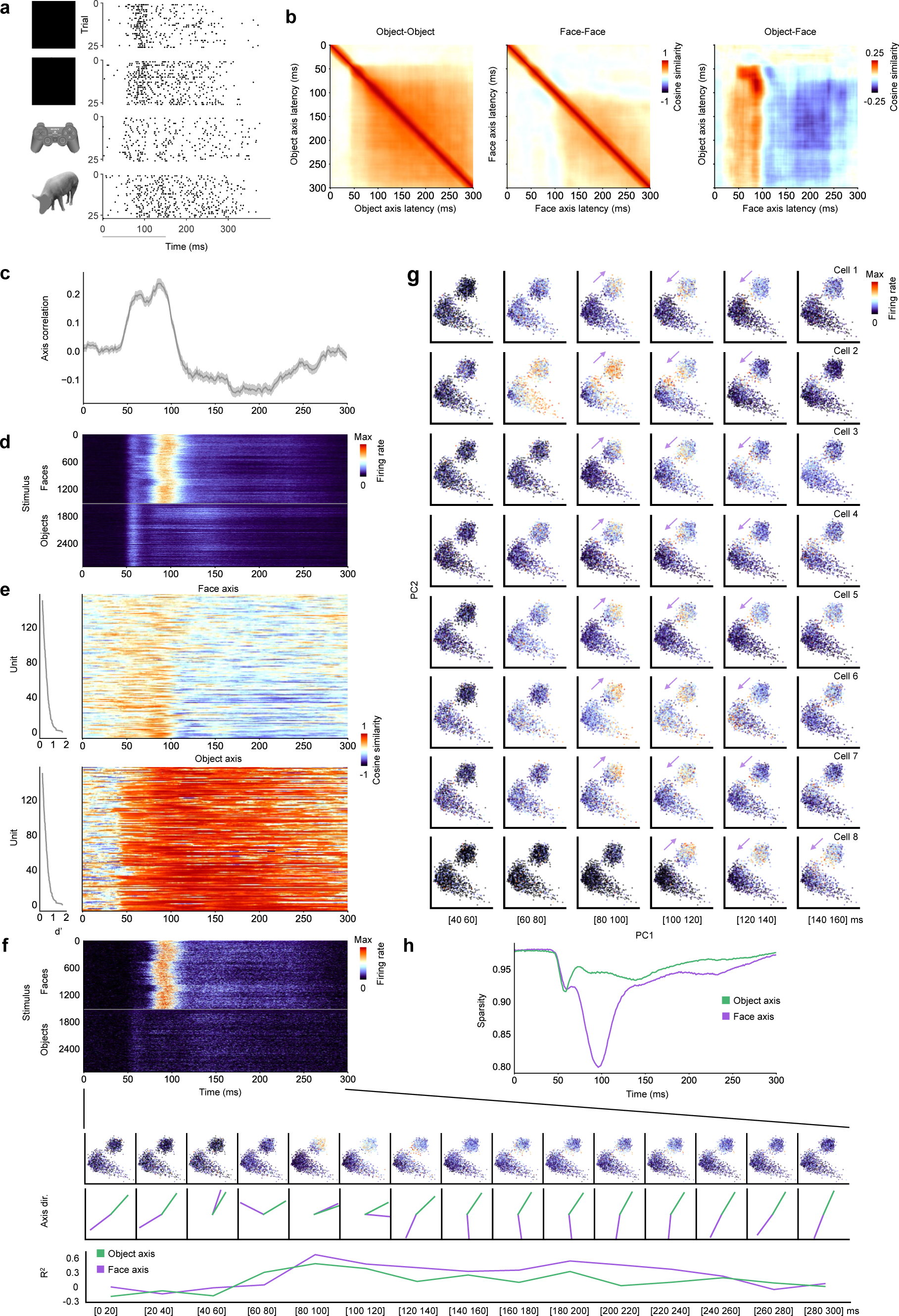
Face and object axes transiently align before diverging. [Note: all human faces in this figure have been redacted due to biorxiv policy on displaying human faces.] **a.** Raster plot of responses of an example ML cell to two face and two non-face stimuli. The gray bar at the bottom indicates stimulus ON time. **b.** From left to right: matrices of mean cosine similarities across the population between (object, object), (face, face), and (face, object) axes (computed using all 60 dimensions) for different pairs of latencies (N = 151 cells; only cells with R^2^ > 0 on test set for object and faces axes were used, see Methods). **c.** Correlation between face and object axes as a function of time (diagonal values in the rightmost similarity matrix in (b)). The shaded region indicates standard error of the mean. **d.** Mean response time course to each face and object stimulus, averaged across cells and trials. **e.** Cosine similarity between the overall object axis for each cell (computed using a time window of 50-220 ms) and its time-varying face (top) and object (bottom) axes, sorted from top to bottom according to face selectivity d’ (left). **f.** Axis change dynamics in a single cell. Top: mean response time course of this cell to each face and object stimulus, averaged across trials. Expanded Row 1: time-resolved scatter plots of face and object stimuli projected onto PC1 and PC2 of object space, color coded by the cell’s response magnitude. Expanded Row 2: time-resolved face and object axis directions computed using a time window of 20 ms. Expanded Row 3: explained variance (R^2^) on test set for object axis (green) and face axis (purple) as a function of time. **g.** Eight example cells showing axis reversal in PC1-PC2 space for faces but not objects. **h.** Response sparsity as a function of time for faces and objects (see Methods).

It is unclear whether such firing rate changes over time reflect *dynamic changes in the neural code for faces and objects.* The axis encoding framework gives us a tool to directly address this. To observe potential axis change dynamics, we next computed face and object axes using a 20 ms sliding window. **Fig. 3b** shows matrices of mean similarities across the population between (object, object), (face, face), and (face, object) axes for different pairs of latencies. The object axis became stable at 60 ms (half-peak time), while the face axis became stable later at 118 ms (**Fig. 3b**, left and middle). Surprisingly, the (face, object) matrix showed a positive correlation very early, in the interval 50-100 ms (**Fig. 3b**, right; **Fig. 3c**). This time period included the time of maximal responses to faces across the population (93 ms, **Fig. 3d**). After 100 ms, the (face-object) axis correlation then became negative (**Fig. 3c**). Thus the face axis becomes transiently aligned to the object axis at short latency before later diverging.

To examine axis dynamics within single cells, we computed the cosine similarity between the overall object axis for each cell (computed using a time window of 50-220 ms) and its time-varying face and object axes (**Fig. 3e**). The face axis showed a strong correlation to the object axis at a short response latency (50-100 ms) but then diverged (**Fig. 3e**, top). The object axis, in contrast, showed high correlation throughout the entire presentation duration (**Fig. 3e**, bottom). The cells in **Fig. 3e** are sorted according to their face selectivity d’; divergence was more prominent for highly face-selective cells.

Inspection of raw responses of an example cell projected onto PC1-PC2 space (**Fig. 3f**) revealed that over time, the face axis did not merely diverge from the object axis, it actually *reversed direction*. In this cell, face and object axes were aligned at 80-100 ms, both pointing in the face quadrant of PC1-PC2 space. Thereafter, the face axis suddenly reversed direction at 120-140 ms. In contrast, the object axis did not change over time.

This pattern of reversal of the face axis in PC1-PC2 of the general object space at ∼100 ms was extremely common across the population. **Fig. 3g** shows the reversal in 8 additional cells. Across the population, 62% of cells that had clearly flipped tuning in PC1-PC2 space (**Extended Data Fig. 8a**; angle ≥ 120°). The reversal was accompanied by sparsening of face cell responses (**Fig. 3h**).

What do the first two components of the object axis, to which face axes are initially aligned but then become reversed, encode? **Extended Data Fig. 8b** shows face and object images projecting maximally onto the extremes of these two components of the mean object axis. Within the face distribution, the images with minimal projection tended to have lower contrast and less distinct eyes. Thus one concrete interpretation of face axis reversal is that face images with minimal projection simply evoked strong responses slightly later; we address and rule out this possibility below (cf. **Extended Data Fig. 12j-l**). Moreover, we emphasize again that the reversal only occurred for faces and thus cannot be simply attributed to a general increased latency to low contrast stimuli.

The fact that face axes *precisely reversed direction* in low dimensions of the general object space just 20-40 ms after they were aligned was a surprise, given our finding using a time-averaged response of a widely divergent face axis distribution relative to the object axis in PC1-PC2 space (**Fig. 2a**). In retrospect, the latter finding was likely due to temporal blurring, illustrating the importance of analyzing IT encoding axes in a time-resolved way.

Cells in face patch AM showed a largely similar pattern of dynamics, but with longer latency overall (**Extended Data Fig. 9**). In particular, in AM the face axis also transiently aligned with the object axis early on (at ∼100 ms) before reversing in low dimensions (interestingly, there was also a weak, brief second period of re-alignment at ∼160 ms not observed in ML). The proportion of cells that had clearly flipped tuning in AM was 57% (**Extended Data Fig. 8a**). The fact that face axis reversal in AM occurred later than in ML suggests that the ML reversal was not due to long-range feedback from AM.

A shadow of these rich dynamics can even be discerned by simply looking at the average responses of cells to the stimuli over time (**Fig. 3d**). At short latency (∼60 ms), there was an initial response which appeared non-specific when averaged across cells (this very short-latency, transient response occurred in 129/151 cells in ML, but not in AM, and was non-, face-, or object-selective depending on the cell). Next, the response to faces became maximal: this is the phase when the face axis is aligned to the object axis. Finally, the face response sparsened: this is the phase when the face axis reversed direction. Objects, in contrast, evoked a sustained response without sparsening, reflecting the fact that the object axis continually pointed towards the face quadrant without change (**Fig. 3e-g**).

Overall, these results reveal temporal dynamics within the face patch population in response to faces but not to non-face objects, particularly the striking phenomenon of reversal of the face axis relative to the object axis in the general object PC1-PC2 space following initial alignment. This dynamic means that while a single axis can explain the initial response of face cells to faces and objects, this is no longer true after the reversal event (thus coding becomes domain specific, cf. **Fig. 1d**).

### Face axis reversal in low dimensions is accompanied by emergence of diverse new tuning to higher dimensions of face space

What is the functional significance of face axis reversal in PC1-PC2 space? In broad terms, reversal should endow cells with a greater dynamic range for representing differences between faces by removing the common response due to face detection. To gain further insight, we trained a recurrent neural network (RNN) to produce a similar reversal in response (see Methods). The weight matrix that emerged showed surprisingly simple structure consistent with lateral inhibition, exhibiting local inhibitory weights and longer range excitatory weights^39^ (**Extended Data Fig. 10**). Thus we hypothesized that axis reversal might be an indicator of lateral inhibition in the biological brain. Since lateral inhibition is a powerful mechanism to sculpt feature tuning^40,41^, we further hypothesized that axis reversal might be concomitant with *emergence of new tuning to facial details,* and this should only occur in response to face stimuli–since object stimuli never triggered axis reversal.

To look for new tuning to face features, we first generated a 60-d *face space* by passing face stimuli through AlexNet and performing PCA on the fc6 representation (cf. **Fig. 1b**). This new AlexNet face space emphasizes differences between face identities rather than between faces and objects and explained 86.4% of variance in the fc6 features of the face stimuli. A plot of face and object axis weights computed in this new space using a short (80-100 ms) and long (120-140 ms) latency time window revealed a strikingly different pattern for faces (**Fig. 4a**, top and middle) compared to objects (**Fig. 4b**, top and middle). For faces, the weights were strongly anti-correlated between short and long time windows in low dimensions, and uncorrelated in higher dimensions, resulting in a distribution of correlations skewed toward negative values when computed over all 60 dimensions (**Fig. 4c**; one sample t-test, t(150)=-7.78, p<1.1*10^-12^). In contrast, for objects, the weights were clearly correlated between the two time windows across all dimensions (**Fig. 4b, c**; one sample t-test, t(150)=16.01, p<2.8*10^-34^). Importantly, we confirmed that axis weights for faces computed using a short and long time window in higher dimensions 6 - 60 were not positively correlated (**Fig. 4d**, one sample t-test, t(150) = −3.13, p = 0.002), in contrast to axis weights for objects (**Fig. 4d**, one-sample t-test, t(150) = 14.56, p < 1.7*10^-30^); thus the changes in tuning were not confined to low dimensions of the face space. Following the drastic change in face axis weights between 80-100 ms and 120-140 ms, the face axis then stabilized (**Fig. 4a**, middle and bottom, **Fig. 4e**; face: one sample t-test, t(150) = 29.3, p < 5.8*10^-64^, object: one sample t-test, t(150) = 19.1, p < 5.5*10^-42^; cf. **Fig. 3b**).

**Figure 4:**
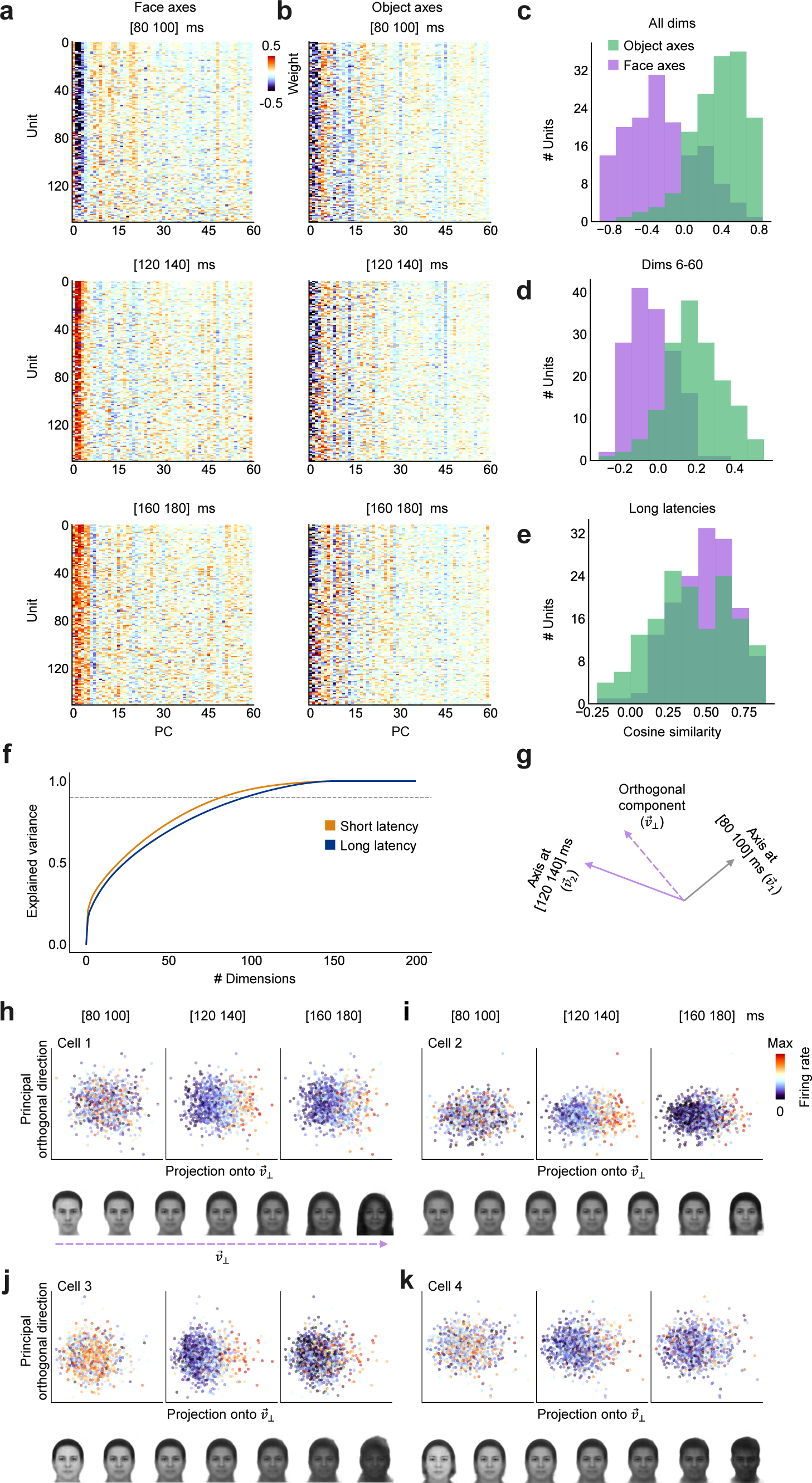
Face axis reversal in low dimensions is accompanied by emergence of diverse new tuning to higher dimensions of face space. [Note: all human faces in this figure were synthesized using a variational autoencoder^58^ and do not depict any real human, in accordance with biorxiv policy on displaying human faces.] **a.** Matrix of face axis weights in AlexNet face space for each cell, computed using a short (80- 100 ms, top), long-early (120-140 ms, middle), and long-late (160-180 ms, bottom) latency window (N = 151 cells; only cells with R^2^ > 0 on test set for object and faces axes were used, see Methods). **b.** Matrix of object axis weights computed at three different latencies; conventions same as (a). **c.** Purple (green) distribution shows cosine similarities across units between face (object) axis weights at short and long-early latency. All dimensions (1-60) were used to compute cosine similarities shown here. **d.** Same as (c), using higher face space dimensions 6-60 to compute cosine similarities for each cell. **e.** Same as (c), showing cosine similarities across units between face (object) axis weights at long-early and long-late latencies (120-140 ms and 160-180 ms), using all 60 dimensions. **f.** Explained variance in z-scored population responses to faces at short (80-100 ms, orange) and long (120-140 ms, blue) latencies, as a function of number of PCs. **g.** Schematic showing projection of a cell’s encoding axis at 120-140 ms (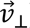) onto the component orthogonal (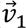) to the cell’s encoding axis at 80-100 ms (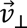). **h-k.** Top: scatter plots of responses to faces projected onto 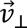 (top) at three different time windows (80-100 ms, 120-140 ms, 140-160 ms) for four example cells. Bottom: faces spanning 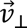 for each cell, sampled at [−4, −2, −1, 0, 1, 2, 4] σ, where σ is the standard deviation of the stimuli along 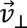 (see Methods).

In **Fig. 4a**, new tuning to many higher dimensions is apparent. This led us to wonder whether we might observe a difference in the dimensionality of the neural state space itself between short (80-100 ms) and long (120-140 ms) latencies, without interpreting responses in terms of tuning to any specific stimulus parametrization. To address this, we performed PCA on raw neural population responses to faces at the two latencies. **Fig. 4f** shows that indeed, more dimensions were required to explain 90% of response variance at long compared to short latency (96 dim for long, 82 dim for short).

The change in axis weights at ∼100 ms implies *emergence of tuning to new dimensions.* To identify these new tuning dimensions, for each cell, we first computed its preferred axis at 80-100 ms (*v̅*_1_) and at 120-140 ms (*v̅*_2_). We then computed the component of *v̅*_2_ orthogonal to *v̅*_1_ (*v̅*_⊥_) (**Fig. 4g**). This component represents a new tuning direction that is orthogonal to the cell’s previous tuning direction. This analysis approach readily revealed tuning to new dimensions emerging around the time of reversal in single cells (**Fig. 4h-k**, top). In each of the four example cells shown, new tuning is apparent along *v̅*_⊥_ at 120-140 ms. We visualized the new tuning directions by generating faces varied along *v̅*_⊥_ (**Fig. 4h-k**, bottom). Each of the four cells showed new tuning to different features, e.g., *v̅*_⊥_ for Cell 1 varied from small inter-eye distance and sharp chin to large inter-eye distance and round chin.

Cells in face patch AM showed a very similar pattern, with reversal in low dimensions and new feature tuning developing in higher dimensions for faces, and no change in tuning for objects (**Extended Data Fig. 11**).

What mechanism underlies the axis change between short and long latencies? Above, we suggested that the switch arises from lateral inhibition, i.e., local recurrent computation. Is it possible that instead, the axis change depends only on a cell’s intrinsic response magnitude to a stimulus without necessitating recurrence? We next investigated and ruled out three “cell-intrinsic” scenarios.

In scenario one, axis dynamics is always accompanied by a high mean response magnitude (**Extended Data Fig. 12a**). To address this, we identified the 100 most effective non-face stimuli as well as the 100 least effective face stimuli (**Extended Data Fig. 12b**). The former evoked a *greater mean response*, averaged over 50-220 ms, than the latter (**Extended Data Fig. 12c**; mean response ratio = 1.29). However, the two types of stimuli evoked very different response dynamics (**Extended Data Fig. 12d**), and comparison of axis weights across different time windows revealed axis change only in response to the face stimuli (**Extended Data Fig. 12e-i**). This shows that the presence of discriminable face features is necessary to trigger axis change, high mean response magnitude is insufficient.

In scenario two, axis change dynamics could be due to delayed responses to weaker stimuli leading to a change in tuning (**Extended Data Fig. 12j**). To address this, we first identified the 50% most and 50% least effective face stimuli for each cell, determined by the mean response in the time window 80-100 ms. We then computed face axes separately using these two groups of stimuli, at both short (80-100 ms) and long (120-140 ms) latency. If axis change dynamics were driven solely by delayed onset of weak stimuli, then one would predict: 1) lack of correlation between axes computed using most and least effective faces at long latency, and 2) positive correlation between axes computed using the most effective faces at short and long latency. Neither of these predictions was supported by the data (**Extended Data Fig. 12k, l**). In particular, the negative correlation that we observed experimentally between axes computed using the most effective faces at short and long latency rules out that possibility that axis reversal is driven solely by responses to non-optimal, low contrast stimuli (**Extended Data Fig. 12l**).

In scenario three, axis change dynamics could be due to a cell-intrinsic increase in firing threshold following a strong transient response, i.e. adaptation (**Extended Data Fig. 12m**). To investigate this possibility, we measured axis tuning of model cells encoding a single axis with and without application of a raised threshold. We found that resulting axes were highly correlated (**Extended Data Fig. 12n**). This contrasts with our actual results (**Fig. 4a-d**), ruling out cell-intrinsic adaptation as the source of the neural code change. Furthermore, in a separate experiment, we presented intact and degraded faces to cells, and found that certain types of degradations consistently led to strong, prolonged responses compared to the response to clear faces (**Extended Data Fig. 12o**). This shows that face cells are not intrinsically wired to always “adapt” following a strong response; instead, the period of high firing can change depending on stimulus context, supporting recurrent mechanisms over cell-intrinsic mechanisms for determining response dynamics.

Together, these three analyses strongly suggest that axis change dynamics cannot be explained by cell-intrinsic changes related to mean firing rate, peak firing rate, or adaptation. As a final control for a non-recurrent mechanism for axis change, we considered whether a “coarse to fine” progression in arrival of visual information to a face patch could explain the change in face axis^42,43^. In the early visual system, information related to low spatial frequency components arises faster than that related to high spatial frequency components^44^. To address the contribution of spatial frequency to axis change, we passed low and high spatial frequency-filtered images separately through AlexNet to generate two different face spaces, one capturing low and one capturing high spatial frequency features (**Extended Data Fig. 13a**). When we compared variance explained by combining high and low frequency features with that explained by only high or only low frequency features, the values across all three feature sets were extremely similar for both the early (80-100 ms) and late (120-140 ms) responses (**Extended Data Fig. 13b**); this indicates that the neural code in the late response is not specially dedicated to representing high spatial frequency features. Indeed, when we repeated the analyses of **Figs. 3** and **4** using either the low or high spatial frequency face space, the pattern of results did not change (**Extended Data Fig. 13c-l**). These analyses argue against an explanation for axis change based on timing differences in arrival of low vs. high spatial frequency information. Most importantly, the fact that axis change occurred only for faces strongly argues against such an explanation, as input timing differences should also affect object responses.

Overall, these results show that for faces, a drastic change in tuning occurs across all dimensions of face space at ∼100 ms, involving reversal in low dimensions and loss/gain of tuning in higher dimensions. We found that the change in tuning was only triggered by stimuli containing facial features (**Extended Data Fig. 12a-i**), and it could not be explained by cell-intrinsic changes in firing rate (**Extended Data Fig. 12j-o**) or by delayed processing of high spatial frequency content (**Extended Data Fig. 13**). Regardless of precise underlying mechanism, the results demonstrate a new form of cooperative computation that challenges the concept that single cells represent fixed features.

### Newly emergent face encoding axes improve face discrimination

What do the newly emergent encoding axes for faces accomplish? Do these changes in tuning to face features improve face discrimination? To quantify the impact of the tuning changes at the population level, we decided to take a population decoding approach, reconstructing faces using responses from different time windows. This approach enables direct visualization of the aggregate effect of tuning changes across the population.

We linearly decoded face features using face responses from three different windows: (1) a short latency window, (2) a long latency window, and (3) a combined latency window; each of the three windows comprised two sub-windows (short: 50-75 ms and 75-100 ms; long: 120-145 and 145- 170 ms; combined: 62-87 ms and 132-157 ms). For each window, we treated responses from the two sub-windows as if they were from distinct cells, enabling the decoder to assign distinct axes to each sub-window and thus leverage axis dynamics (**Fig. 5a**). We used responses from each window to decode face features according to the equation 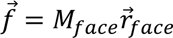 and then passed the decoded features 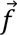 into a variational autoencoder trained to generate faces from latent features (**Fig. 5a**).

**Figure 5:**
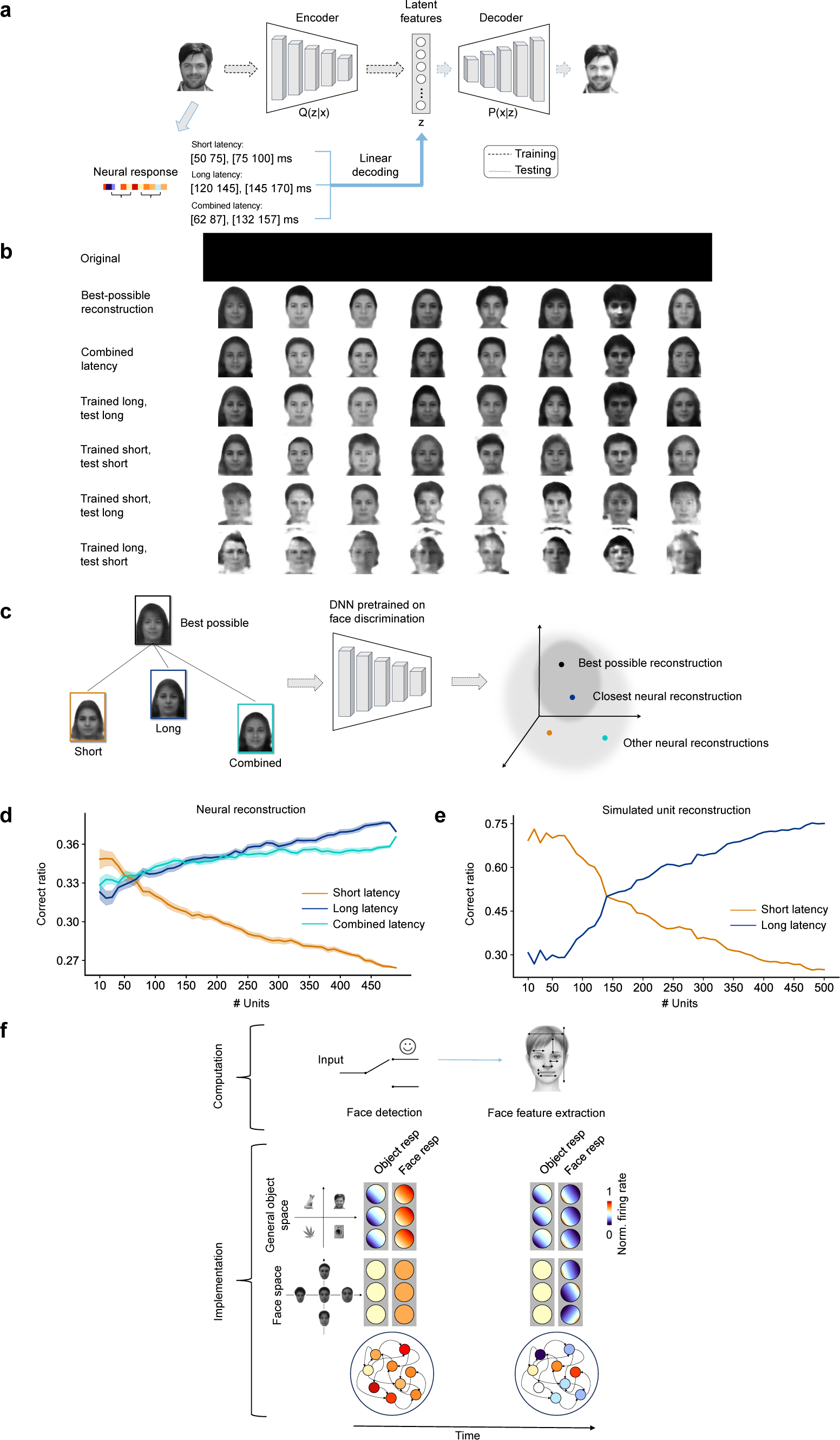
Newly emergent face encoding axes improve face discrimination. [Note: all human faces in this figure were synthesized using either a diffusion model^57^ (panel a) or a variational autoencoder^58^ (panels b, c); they are not photographs. The faces in the first row of panel b have been redacted. Thus all images in this figure are in accordance with biorxiv policy on displaying human faces.] **a.** Schematic of reconstruction pipeline. To reconstruct faces from neural activity, we first trained a probabilistic generative model with an encoder mapping input images to 512 latent features and a decoder projecting the features to the original inputs (see Methods). We then linearly decoded the latent features using neural responses from three different time windows to reconstruct the face. To perform reconstructions, we pooled activity from ML (N = 332 cells) and AM (N = 154 cells); AM time windows were delayed by 20 ms (see Methods). The three windows each comprised two sub-windows to make resulting comparisons fair (i.e., combined and contiguous windows both had two degrees of freedom in axis choice per cell). **b.** Row 1: 8 example faces. Row 2: best possible reconstructions using ground truth feature representations of each face. Row 3: faces reconstructed from neural activity in the combined latency time window, using a decoder trained on combined latency responses. Row 4: Same as Row 3, trained on long latency responses and tested on long latency response. Row 5: Same as Row 3, trained on short latency responses and tested on short latency response. Row 6: Same as Row 3, trained on short latency responses and tested on long latency responses. Row 7: Same as Row 3, trained on long latency responses and tested on short latency responses. **c.** Schematic showing the use of a DNN trained on face discrimination to test identification performance for reconstructed faces. A face was considered identified correctly if it was the closest face in the DNN’s face space to the optimally reconstructed face. **d.** Face identification performance on faces reconstructed from neural activity in different time windows as a function of # cells, following scheme in (c). The shaded area denotes standard error of the mean. **e.** Same as (d), computed using a model in which short latency responses had consistent tuning across units to a small (5/60) number of dimensions while long latency responses had varied tuning across units to a large (60/60) number of dimensions (see Methods). **f.** Summary schematic illustrating implementation of a face detection gate through recurrent dynamics. In the implementation scheme: the three disks in the gray boxes denote three different cells; an initial stage of filtering for face-diagnostic features is achieved via encoding axes for both faces and objects that are aligned and pointing towards the face quadrant (left top). At ∼100 ms, the encoding axis for faces undergoes a reversal in low dimensions of the object space (right top). Simultaneously, new encoding axes develop in the face space (right bottom).

For the long and combined windows, this approach gave reasonable reconstructions compared to the best possible reconstruction (i.e., the reconstruction from veridical face features, see Methods) (**Fig. 5b**, rows 3 and 4). Reconstructions were less accurate using the short window (**Fig. 5b**, row 5). When we attempted to perform cross-window decoding by training a decoder using short (long) window responses and then applying the decoder to long (short) window responses, face reconstruction completely failed (**Fig. 5b**, rows 6 and 7). This underscores the complete change in neural code occurring at ∼100 ms (**cf. Fig 4**).

We quantified reconstruction quality by identifying the reconstructed faces using a state-of-the-art DNN^45^ pre-trained to discriminate different celebrities in the VGGFace-2 dataset^46^. We compared faces reconstructed using short, long, and combined windows, asking which reconstruction was closest to the best possible reconstruction in the VGGFace-2 DNN space (**Fig. 5c**). For small numbers of cells, the short latency response performed best (**Fig. 5d**, orange curve). However, as cell number increased, the long and combined responses outperformed the short latency response (**Fig. 5d**, blue and cyan curves). We also compared face reconstruction accuracy by quantifying how well we could identify reconstructed faces among distractors; these results confirmed that long and combined windows outperformed the short window (**Extended Data Fig. 14**).

The pattern of recognition performance in **Fig. 5d** exactly matches what one would expect if the short latency response were specialized for face detection and the long latency response for face discrimination. For face detection, cells only need to represent a set of dimensions diagnostic of all faces. Tuning to a smaller number of dimensions leads to increased robustness due to redundancy in tuning. In contrast, for face discrimination, cells need to represent a larger set of dimensions enabling fine differentiation. Simulation of this tradeoff (redundancy vs. diversity) produced the same pattern of responses as we observed in the actual cell population (**Fig. 5e**).

Overall, these results show that the drastic changes in feature tuning occurring at ∼100 ms have a functional consequence, markedly improving the ability of the neural population to perform fine face discrimination.

## Discussion

In this paper, we resolve a longstanding debate about whether the coding mechanism used by IT cortex is domain specific^17–19^ or domain general^20–22^. We show that face cells initially adopt a domain-general code, but following a rapid, concerted population event lasting < 20 ms, the neural code transforms into a domain specific one. Furthermore, our results suggest that the initial domain-general phase is performing face detection, while the later domain-specific phase is performing face discrimination–providing a concrete neural implementation of the long-hypothesized face detection gate for explaining domain-specific face processing^23^ (**Fig. 5f**). At short latency, cells use a common encoding axis for faces and objects that points towards faces in a DNN-derived feature space, optimally subserving face detection. In contrast, at longer latency, the same cell population switches to *face-specific* axes to encode faces. This phase is marked by reversal of the face axis in low dimensions of the object space (i.e., cells are *no longer detecting faces*), sparsening of responses, and emergence of tuning to new facial feature dimensions leading to improved face discrimination.

We believe these findings are fundamentally important for three major reasons. First, the findings reveal a new form of neural computation. Our findings show that the tuning of cells in macaque face patches is not fixed but can change over an extremely short timespan (within 20 ms, cf. **Fig. 3f-h**). Note we are not simply claiming that representation of different features occurs with different latencies, which would be unsurprising given the multiple synaptic paths a stimulus can take to reach IT cortex. Rather, what we have found is a wholesale switch in neural code from object axes mediating face detection to face axes mediating face discrimination; moreover, this switch is *stimulus-gated* and hence *dynamically controllable*. In response to a non-face object, the switch in neural code never occurs, and cells across the face patch population persist in using object axes for the entire stimulus duration.

Several previous studies have observed temporal dynamics in IT visual representation. Sugase et al. reported that information about stimulus category (face versus object) rises ∼50 ms earlier in IT cells than information about stimulus identity^47^. Tsao et al. came to a similar finding, comparing time courses for decoding face/object category versus individual face identity in face patch ML^15^. Our present findings reveal that a *change in neural code* occurs concomitant to the improved decoding performance for face identity. This is not implied by the previous findings. For cells using a single, fixed encoding axis, category information would also be expected to rise earlier than identity information, simply because categorization can rely on larger image differences that produce larger neural signal differences and hence require less temporal integration to decode.

Brincat and Connor found tuning to multipart configurations arises later in IT cortex compared to tuning to individual parts^48^. This finding is also consistent with our results, but we emphasize again that our discovery is distinct from finding different temporal dynamics for extracting different types of features. What is especially novel about the dynamic representational mechanism we have uncovered is that *the dynamics are controllable by the category of the stimulus itself.* In response to an object, face patches persist in using object axes. Only in response to a face does the neural code rapidly switch to face-specific axes.

Our results challenge a prevailing view that “core object recognition,” the ability to rapidly (in < 200 ms) recognize objects despite substantial appearance variation, can be accounted for by largely feedforward processes^37^. Invariant recognition of clear, isolated faces in the absence of any background clutter inarguably falls within the realm of core object recognition. Our results show that this process is always accompanied by a rapid switch in neural code following stimulus presentation in the key brain structures mediating face recognition^30^.

We believe the switch in neural code for faces must be implemented by some form of dynamic computation. One possible scheme would be a feedforward network with different modules, one that computes the object axis response, one that computes the face axis response, which both sum to the recorded cell, and one early module that detects whether something is a face or not and completely shuts down the face or object axis module respectively. Note that to account for the delay in onset of the face-specific axis, different delay lines from each module would be required, which is not part of a standard DNN. Alternatively, since we know that IT cortex harbors rich local recurrent connections^49^ and receives feedback from multiple areas^50^, it is possible that a dynamic switch in encoding axis could be mediated by these local and long-range recurrent connections^5,51^. We favor the latter possibility, especially given the results of our RNN model showing how a simple form of lateral inhibition can lead to axis reversal (**Extended Data Fig. 10**). Regardless, our findings show that core object recognition cannot be explained by classic feedforward mechanisms.

The second major conceptual advance of this study is to provide a new mechanistic framework for understanding a large body of behavioral and neuropsychological work on face perception. Behavioral studies indicate much better discrimination of face parts when presented in the context of a whole face^24,25^ (cf. **Fig. 1a**), and neuropsychological case studies point to a double dissociation between face and object recognition circuits^52^. At a high level, both of these observations can be accounted for by a face detection gate^23^. Our findings suggest that face patches implement a face detection gate in a remarkably literal sense, as an initial stage of filtering for face-diagnostic features prior to detailed analysis of facial identity (**Fig. 5f**). Intuitively, the picture arising from our results is that holistic processing constitutes a type of “zoomed-in” vision: only stimuli which pass the face detection gate can be perceived with this special form of vision. Our findings also resonate with the context frame theory of visual scene processing^42^, according to which an early wave of processing encodes coarse, context-based predictions–a context frame–while a later wave encodes detailed recognition.

Finally, our results provide a concrete answer to a fundamental question that has stubbornly eluded the field of face perception: *what is a face?* While cells in face patches invariably respond more strongly to faces than to, say, scissors, one can often find cells that respond strongly to a pair of small disks or a simple oval^53^. Are these faces too? Remarkably, our results suggest that an unambiguous answer may exist: we hypothesize that a face patch considers a stimulus a face if the stimulus triggers the *state change* leading to axis reversal and concomitant new feature tuning. Indeed, we found that animal bodies with grayed-out heads evoked strong responses across the face cell population^54^, but these stimuli did not trigger the state change associated with actual faces. Instead, they elicited a prolonged form of the initial detection phase, i.e., they were treated by the face patch as object stimuli.

Since the advent of AlexNet, there has been intense interest in using DNN models to study high-level vision^2,4,6,22,55^. In particular, Vinken et al. performed experiments similar to ours, recording responses of macaque face cells to faces and objects, and computed encoding models for each using a common DNN representation^22^. However, they came to a diametrically opposite conclusion, namely, that the computation performed by face cells is domain general and can be accounted for by a single, generic encoding model. Like us, they also found higher accuracy for explaining responses to faces using a face-specific encoding model compared to an object-encoding model, but they concluded that this was a consequence of out-of-distribution generalization error. We believe two critical pieces of data refute their interpretation. First, the face axis is not only different, but becomes diametrically reversed relative to the object axis in low dimensions (**Fig. 3f, g, Extended Data Fig. 8**). Second, at very short latencies, the face axis *is significantly correlated* to the object axis (**Fig. 3b, c, e-g**); thus the observed *lack of correlation* at longer latencies is a real signal. The critical step to coming to our findings is analyzing face and object axes in a time-resolved manner.

An objection that might be raised to our conclusions is that we only tested AlexNet fc6 feature representations. Couldn’t some other more sophisticated network harbor more nonlinear representations that would make the difference between face and object axes disappear? In effect, such a network would harbor single axes expressing “if face then use face axis; if object then use object axis.” In our opinion, two linear models are more parsimonious than one highly nonlinear blackbox model.

In a remarkably prescient paper, Keiji Tanaka speculated on the function of cortical columns^56^. He noted that the anatomical organization of clustered functional activity with extensive horizontal arborizations enables cells to operate either in “pooling mode,” where they encode the common feature preference across cells, or in “differential amplifier mode,” where they encode fine differences in feature preferences. Our results show that the face patch network switches between these two modes each and every time a face is perceived. Given that IT cortex contains multiple shape and color networks organized similarly to the face patch network^6,11^, we believe that the present findings will likely generalize to other parts of IT, and object detection and discrimination via dynamic feature axes may be a general computational principle of IT cortex. More broadly, the mechanism for switching attention from the whole (face detection) to the parts (detailed face discrimination) may constitute a fundamental circuit for perceiving compositionality. Cognition is at heart a process of dissecting hierarchical structure, and axis change dynamics may provide a general mechanism to attend to different levels of hierarchy—whether in vision, language, or logic.

## Methods

Two male rhesus macaques (Macaca mulatta) were used in this study. All procedures conformed to local and US National Institutes of Health guidelines, including the US National Institutes of Health Guide for Care and Use of Laboratory Animals. All experiments were performed with the approval of the UC Berkeley Institutional Animal Care and Use Committee.

### Visual stimuli

#### Face patch localizer

The fMRI localizer stimulus contained 5 types of blocks, consisting of images of faces, hands, technological objects, vegetables/fruits, and bodies. Face blocks were presented in alternation with non-face blocks. Each block lasted 24 s (each image lasted 500 ms). In each run, the face block was repeated four times and each of the non-face blocks was shown once. A block of grid-scrambled noise patterns was presented between each stimulus block and at the beginning and end of each run. Each run lasted 408 seconds.

### Stimuli for electrophysiology experiments

#### 1) Human faces

2000 frontal views of faces, as in ^1^, were acquired from various face databases: FERET^2,3^, CVL^4^, MR2^5^, Chicago^6^, CelebA^7^, FEI (fei.edu.br/∼cet/facedatabase.html), PICS (pics.stir.ac.uk), Caltech faces 1999, Essex (Face Recognition Data, University of Essex, UK; http://cswww.essex.ac.uk/mv/allfaces/faces95.html), and MUCT (www.milbo.org/muct). The faces were aligned as in Ref. ^1^, using an open-source face aligner (github.com/jrosebr1/imutils).

#### 2) Non-face objects

A set of object images consisting of 1,392 different images of isolated objects, previously used in Ref. ^8^, were used.

In monkey A, we presented a subset of 1525 faces and 1392 objects. In monkey J, we presented all 2000 faces and 1392 objects (a smaller number of faces were presented to monkey A in order to increase the number of repetitions per stimulus; analyses in monkey J used the same subset of 1525 faces that were presented to monkey A). For all experiments in monkey A, we showed the human faces and objects in separate blocks. Within each block, we showed a training set (a random subset of 1425 faces or 1292 objects) for 1 repetition, followed by the test set (the remaining subset of 100 faces or 100 objects) for 3 repetitions. The two blocks were presented repeatedly until enough repetitions were acquired (around 10 repetitions per training image and 30 repetitions per test image). For all experiments in monkey J, images from different categories were not separated in blocks; instead, faces and objects were randomly drawn and interleaved. All images were rendered in grayscale and subtended 3.9° x 3.9°.

In addition, we presented a screening stimulus set consisting of 6 object categories (faces, bodies, hands, gadgets, fruits, and scrambled patterns) with 16 identities each (**Extended Data Fig. 3**).

### Behavioral task

For electrophysiology and behavior experiments, monkeys were head fixed and passively viewed a screen in a dark room. Stimuli were presented on an LCD monitor (Asus ROG Swift PG43UQ). Screen size covered 26.0° x 43.9°. Gaze position was monitored using an infrared camera eye tracking system (Eyelink) sampled at 1000 Hz.

#### Passive fixation task

All monkeys performed a passive fixation task for both fMRI scanning and electrophysiological recording. Juice reward was delivered every 2-4 s in exchange for monkeys maintaining fixation on a small spot (0.2° diameter).

### MRI scanning and analysis

Subjects were scanned in a 3T PRISMA (Siemens, Munich, Germany) magnet. 1) Anatomical scans were performed using a single loop coil at isotropic 0.5 mm resolution. 2) Functional scans were performed using a custom eight-channel coil (MGH) at isotropic 1 mm resolution, while subjects performed a passive fixation task. Contrast agent (Molday ION) was injected to improve signal/noise ratio. Further details about the scanning protocol can be found in ^9^.

#### MRI Data Analysis

Analysis of functional volumes was performed using the FreeSurfer Functional Analysis Stream^10^. Volumes were corrected for motion and undistorted based on acquired field map. Runs in which the norm of the residuals of a quadratic fit of displacement during the run exceeded 5 mm and the maximum displacement exceeded 0.55 mm were discarded. The resulting data were analyzed using a standard general linear model. The face contrast was computed by the average of all face blocks compared to the average of all non-face blocks.

### Electrophysiological recording

We used Neuropixels 1.0 NHP probes^11^ (45 mm long probes with 4416 contacts along the shaft, of which 384 are selectable at any time) to perform electrophysiology targeted to face patches ML and AM. All Neuropixels data was acquired using SpikeGLX and OpenEphys acquisition software and spike sorted using Kilosort3^12,13^ with the threshold parameter set to (10, 4). Both cells designated good units and MUA by Kilosort3 were used for analyses. For better alignment of the guide tube and the probe, we developed a custom insertion system composed of a linear rail bearing and 3D printed fixture enabling a precise insertion trajectory. Patches were initially targeted with single tungsten electrodes prior to Neuropixels recordings, following methods for MRI-guided electrophysiology described in ^14^.

We recorded data from monkey A in 3 sessions (ML: 563, 331, 364 units; AM: 248, 440, 496 units) and monkey J in 3 sessions (ML: 477, 429, 467 units). Data across all six sessions was consistent. Data shown in **Figs. 1-5** was from one session in monkey A; data shown in **Extended Data Fig. 2** was from one session in monkey J.

### Data analysis

Only visually-responsive cells were included for analysis. To determine visual responsiveness, a two-sided t-test was performed comparing activity at −50-0 ms to that at 50-300 ms after stimulus onset. Cells with p-value < 0.05 were included. Wherever further cell selection was performed (e.g., to cull cells whose activity could be well explained by an axis model, as determined by R^2^), this is indicated in the relevant section of the manuscript.

Here, we summarize these inclusion criteria: In **Figs. 2-4**, we analyzed the same subset of cells that satisfied all of the following criteria: significant visual responsiveness to our main stimulus set consisting of 1525 faces and 1392 objects; nonzero response variance across stimuli; peak *d*’ ≥ 0.2 in 80-140 ms (see “Face selectivity index” below); and the presence of positive R^2^ on a held-out test set for both face and object axes, in both the general object space and the face space (see “AlexNet general object space and face space” below), between 80-140 ms. This resulted in 151/563 units in ML of monkey A, 76/248 units in AM of monkey A, and 131/467 units in ML of monkey J. In **Fig. 5**, in order to include a larger population of cells to improve face reconstruction, we selected cells that responded significantly differently to faces and to non-face objects using a different inclusion criterion: a two-sided t-test was performed comparing each cell’s activity to all faces in the screening stimulus set (consisting of 6 categories of images) to that to all objects in the screening set. Cells with p-value < 0.05 were included.

### Face selectivity index

A face selectivity *d*’ was defined for each cell and each 1 ms time bin as:

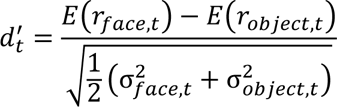

Where *E*(*r*_t_) and 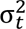 are the mean and variance of responses of the cell over stimuli at time *t*, respectively. We further calculated the peak *d*^′^ of a 20 ms sliding window between 80-140 ms (corresponding to the time interval in which we observed the main face-selective responses in raw time courses, cf. Fig. 3d):

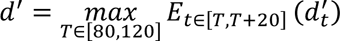

We then selected for cells with peak *d*^′^ ≥ 0.2.

In Extended Data Fig. 3, face selectivity index (FSI) was defined for each cell as:

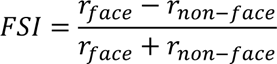

where *r* is the average neuronal response in a 50-220 ms window after stimulus onset.

### Average response profile

Mean responses of each cell to each stimulus were computed in a 50-220 ms window after stimulus onset. Responses were then normalized for each cell to the range [0 1], where the minimum response was assigned 0 and maximum was assigned 1.

AlexNet general object space and face space

We used AlexNet^15^ to embed stimulus images into a high-dimensional latent space. Specifically, images were passed through a pre-trained version of the model in MATLAB (Mathworks), and 4096-dimensional features were extracted from layer fc6. To further reduce dimensionality, PCA was performed. We built two feature spaces using this approach, a 60-d general object space and a 60-d face space.

To build the 60-d general object space, we performed PCA on fc6 responses to a set of 100 face images (randomly selected from the 2000 faces in our face database) and 1292 object images. We normalized each dimension so that the projection of all stimuli along the dimension had mean 0 and standard deviation 1. This general object space was used for the analyses shown in **Figs. 2, 3**, and related **Extended Data Figs. 2a-j, 4-9**. For visualization purposes, we also used the 2D subspace consisting of the first two PCs of this space, as in Ref. ^8^.

To build a 60-d face space capturing variation in face-specific features, we performed PCA on fc6 responses to a set of 1425 face images. This face space was used for analyses shown in **Fig. 4** and related **Extended Data Figs. 2k-q and 11-13**.

### Preferred axis of cells

The preferred axis of each cell was computed using linear regression as follows:

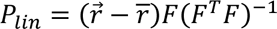

where *r̅* is 1 × *n* vector of the firing rate response to a set of *n* face stimuli, *r̅* is the mean firing rate, and *F* is a *n* × *d* matrix, where each row consists of the *d* parameters representing each image in the feature space. As mentioned above in “Stimuli for electrophysiology experiments,” we split the full image set into a train and test set, and linear regression models were trained on the training set and tested on the held-out test set. R^2^ is shown for the held-out test set for all figures.

### Stimulus distribution control

To evaluate whether the shape of our stimulus distribution had any effect on the axis directions (Extended Data Fig. 5), we fit multivariate Gaussian distributions to the face and object training sets, and then identified subsets of 500 faces and 500 objects with the highest probability density under the face and object Gaussian distributions, respectively.

### Cross prediction between face and object axes

To evaluate how well the face axis generalizes to object responses and vice versa, we trained a linear regression model with the training set for faces (objects), and then tested on the test set for objects (faces) (Fig. 2c, Extended Data Fig. 2c, 6c, 7c).

### Normalized face-object axis correlation

To calculate the correlation between face and object axis of each cell while taking into account the OOD effect, we first measured how well the face or object axis correlated to itself when we calculated the axis using two different subsets of images. We used the mean of face subset-face subset correlation and object subset-object subset correlation of a cell as the best-possible correlation with the OOD effect present. Then we calculated the correlation between the face and object axis of each cell, and took the ratio between this raw correlation and the best-possible correlation computed earlier as the normalized face-object axis correlation (Extended Data Figs. 2d, 6d, 7d).

### Artificial unit comparison

To observe how a unit with a single axis across both face and object space would behave, we identified artificial face-selective units from the AlexNet fc6 layer and redid our analyses for real neural units on them (Extended Data Fig. 7). Since the general object space was built by PCA from the activities of AlexNet fc6 layer unit activities, a unit in the same layer should respond linearly to the features in the general object space, thus providing a model neuron with a single encoding axis. Since there are features not perfectly explained by the feature space, the artificial units should preserve effects from out-of-distribution generalization. We first identified face-selective AlexNet fc6 units by calculating FSI for each unit using its response to the screening stimulus set. We then repeated the analyses of Fig. 2a-c using responses of the 126 most-face selective units (FSI ranging from 0.23 to 0.46).

### Time varying axis analysis

We fit axes to average neural responses in 20 ms windows over the trial duration of 0-300 ms after stimulus onset (Figs. 3, 4 and related Extended Data Figures). To quantify alignment between the face and object axes at different latencies, we computed cosine similarity between the face axis at each latency and the trial-wide object axis computed across 50-220 ms (Fig. 3e, Extended Data Figs. 2h, 9d).

### Identifying cells with an ultra-short latency response

Units were tested for the presence of a response between 50-60 ms as follows: a unit was defined as having an ultra-short latency response if the mean response between 50-60 ms was significantly greater than the mean response between 40-50 ms, assessed by one-sided t-test p-value < 0.05 with Bonferroni correction.

### Population response sparsity

To characterize the sparsity of face cell population responses to faces and objects, we computed a modified Treves-Rolls population sparseness^16^. Specifically, we calculated

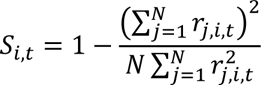

where *N* is the number of neurons and *r*_*o*,*l*,*t*_ is the response of neuron *j* to stimulus *i* at time *t*, for all stimuli and times. To compare population response sparsity to faces and objects, we took the mean over all face and all object stimuli, respectively, at each time point.

### RNN training

We trained a recurrent neural network with 100 units and dynamics at each timestep described by

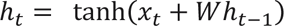

where *h*_*t*_ is the vector of unit activations at time *t*, *w* is the weight matrix, and *x*_*t*_ is the external input at time *t*. During training, the network was run for two timesteps with the same random linear gradient provided as input at both timesteps, and the mean squared error between the output at time 2 and the reversed input gradient was used as the loss. The network was trained on 10,000 iterations, each a random linear gradient, using gradient descent and the Adam optimizer with learning rate 0.001.

### Computing axis correlation between short and long latencies

To quantify changes in face and object axes (Fig. 4c-e and related Extended Data Figures), we computed the dot product between the face axes at three different latencies (80-100 ms, 120- 140 ms, 160-180 ms), and similarly for object axes. We chose to focus on these three time windows because they straddled the peak population response of face cells (Fig. 3d); we skipped 100-120 ms since for many cells, this is when the axis reversal in low components occurred (cf. Fig 3g). We note that individual cells sometimes showed salient changes in response to objects over time; our analyses and conclusions are intended to capture population-wide changes in tuning.

To account for different latencies in responses of different units to faces and objects (e.g., see example cell in Fig. 3f), for each unit, we took the short latency to be the first 20 ms window in which the mean response of that unit was at least 2 standard deviations above the mean baseline (−25−25 ms) response of that unit to faces and objects respectively.

### Identifying new tuning directions for each cell

To identify new tuning directions that emerge at long latency (Fig. 4g-k and related Extended Data Figures), we compared the face axes of each cell at 80-100 ms, denoted as 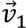, to the axes at 120-140 ms, denoted as 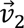 (AM time windows were delayed by 20 ms). We decomposed 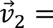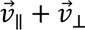, where 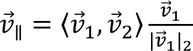 is the component of 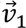 parallel or antiparallel to 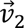, and 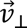, the component of 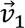 orthogonal to 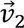, is given by 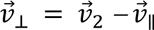 For visualizations in Fig. 4h-k, we computed the principal orthogonal direction for each 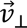. Specifically, for each stimulus with embedding 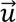 in the face space, we orthogonalized 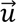 with respect to 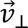 by computing 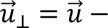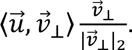. Then, we performed PCA over all 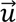 and took the first PC as the principal orthogonal direction to 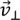.

### Generating low and high spatial frequency-filtered images

Each image from our original set of 1525 face images was first convolved with a Gaussian filter (σ = 0.044°) to generate a low spatial frequency-filtered image. This was then subtracted from the original to compute a high spatial frequency image (Extended Data Fig. 13).

### Face reconstruction

To reconstruct faces from neural activity (Fig. 5, Extended Data Fig. 2q, r, Extended Data Fig. 14), we leveraged a probabilistic generative model^17^. The model followed the design of variational autoencoders^18^, with an encoder mapping the input images (resized to 128×128) into latent features defined with a variational distribution and a decoder projecting the features to the original inputs. The encoder and decoder each consisted of five convolutional layers, and the latent features were set to 512 dimensions. A regularizer was used to minimize discrepancies between the latent distribution and a prior Gaussian distribution, which better supports data on low-dimensional manifolds. We also included an additional objective to align the latent features (after linear projection) with fc6 features from AlexNet, to ensure the latent space described similar features as the general object space. We trained the generative model on 1900 faces and 1292 objects and validated with 100 images each.

To compare the reconstructed faces from neural responses at short and long latency, we used averaged population responses from various time windows. For ML cells, short latency responses were averaged from 50-75 ms and 75-100 ms, long latency from 120-145 ms and 145-170 ms. We also used a combined window composed of sub-windows from 62-87 ms and 132-157 ms. Note that the three windows each spanned exactly the same duration of time. AM time windows were delayed by 20 ms.

For reconstruction, we linearly decoded a 512-d feature vector from neural responses to each image using responses from the short, long, and combined time windows (thus three separate linear decoders were trained). For each of the three windows, we treated responses from the two sub-windows as if they were from distinct cells, enabling the decoder to assign distinct axes to each sub-window. We then fed the latent features into the decoder of the probabilistic generative model to reconstruct the face. For learning linear decoders of neural activity, we trained on 1425 face images (a subset of the 1900 images used for generative model training) and tested on the remaining 100.

We also passed each original face image through the encoder and decoder of the generative model directly to obtain a best possible reconstruction of each face, allowing us to separate fine face feature loss within the model itself from decoding inaccuracy due to the neural signal.

### Visualizing new tuning directions

To visualize new tuning directions of cells (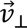, **Fig. 4h-k**), for each cell we reconstructed a series of faces whose features gradually varied along 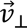, leveraging the probabilistic generative model described above (“Face reconstruction”). Since 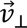 lies in our 60-d face space, we trained a linear transformation matrix to transform features from this 60-d face space to the generative model’s latent space (512-d). We first normalized each cell’s axis to unit length and then sampled seven locations along it, at projection length of [−4, −2, −1, 0, 1, 2, 4] σ, where σ is the standard deviation of the projection of our 1425 original faces onto 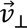. This yielded seven 60-d feature vectors, which we then transformed to 512-d. Finally, we fed these 512-d features into the generative model to generate the reconstructed faces.

### Using a DNN trained on face recognition to test face identification performance

To quantify the goodness of reconstruction (beyond human visual evaluation) (Fig. 5c, d), we leveraged a DNN pre-trained to discriminate faces from the VGGFace-2 dataset^19^, whose feature space provides an objective metric for evaluating similarity between faces. We embedded all our neural-reconstructed faces and best possible reconstructed faces into the feature space of this DNN and then calculated the Euclidian distance between each face’s neural reconstruction and its best possible counterpart.

To compare the goodness of reconstruction among the three time windows, we used two methods. In **Fig. 5c, d**, we compared the distance of reconstructions from the three time windows to the best possible reconstruction for each face. The closest face among the 3 was considered “correct.” We repeated this recognition task for all 1525 faces. We further repeated this whole process using only a subset of the neurons to investigate how number of cells affects the reconstruction quality for different time windows. We repeated this recognition task 30 times by randomly selecting the subset of cells used for the reconstruction. In **Extended Data Fig. 14**, we randomly selected a varying number of distractor faces from our database of 1525 face images and reconstructed these faces from neural data. If the closest face was the target, we considered the decoding “correct.” We repeated this recognition task for 500 targets drawn 30 times and up to 1500 distractors.

### Simulating results of face reconstruction analyses

We simulated a neural population with a short latency response tuned to a smaller set of face dimensions and a long latency response tuned to a larger set of face dimensions, in order to test our hypothesis that the results of Fig. 5d might be explained by a tradeoff between redundancy and diversity.

Simulated neuron responses were created by linearly combining different dimensions of the latent space features of the DNN that was trained on VGGFace-2. We first performed PCA on this DNN’s latent space to get a 60-d feature for each image. Then, for simulating short latency responses of each artificial neuron, we linearly combined a subset of random dimensions from the first 5 dimensions of the features and applied Gaussian noise. For long latency responses, we linearly combined a subset of random dimensions from the full 60 dimensions and applied Gaussian noise. These simulated neurons were then treated as real neurons and underwent the face reconstruction and recognition tasks described above.

## Data availability

All data are available from the lead corresponding author upon reasonable request.

## Code availability

All code are available from the lead corresponding author upon reasonable request.

## Acknowledgments

We thank C. Sohn for animal care support, members of the Tsao laboratory, P. Bao, X. Dai, J. Gallant, J. Gao, S. Kornblith, Y. Ma, F. Tong, and A. Tsao for discussions and critical comments, and P. Bao and L. She for technical advice.

## Author contributions

YS and DYT conceived the project and designed the experiments, YS, JKH, and FFL collected the data, and YS and DB analyzed the data, with help from SC. YS, DB, and DYT interpreted the data. YS, DB, JKH, and DYT wrote the paper, with feedback from FFL and SC.

## Extended Data Figures

**Extended Data Fig. 1:**
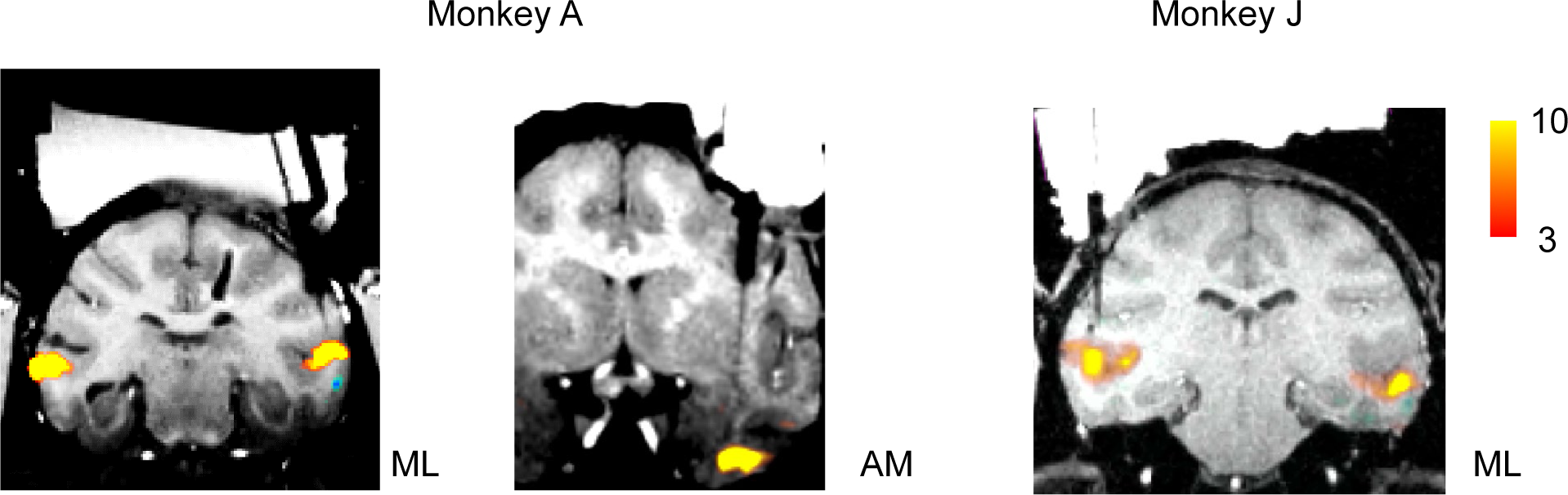
Coronal slices showing the electrode targeting three recording sites from two monkeys. Left: tungsten probe targeting ML in monkey A. Middle: Neuropixels probe targeting AM in monkey A. Right: tungsten probe targeting ML in monkey J. Activations for the contrast faces versus objects are shown, at uncorrected p values in −log10.

**Extended Data Fig. 2:**
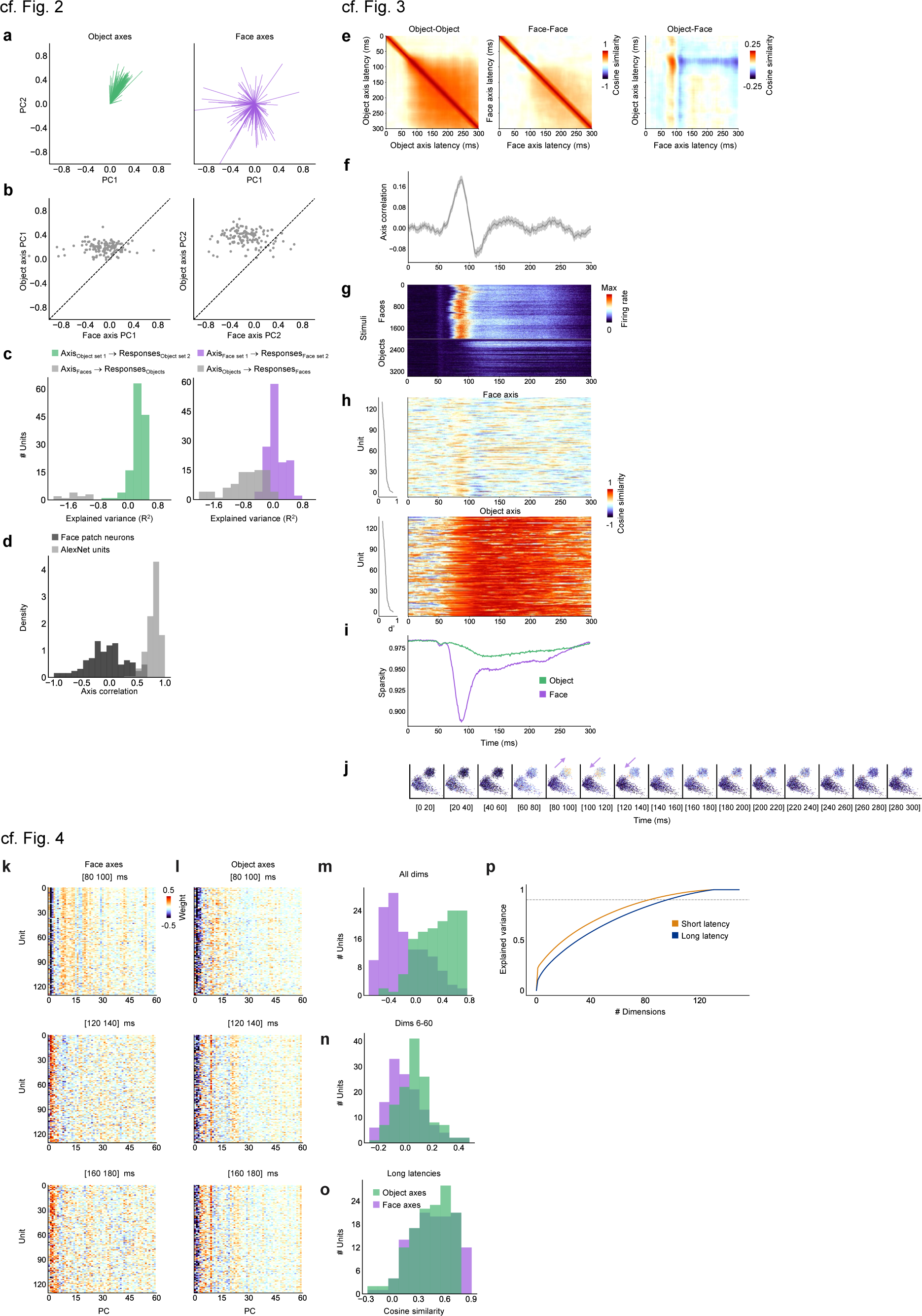
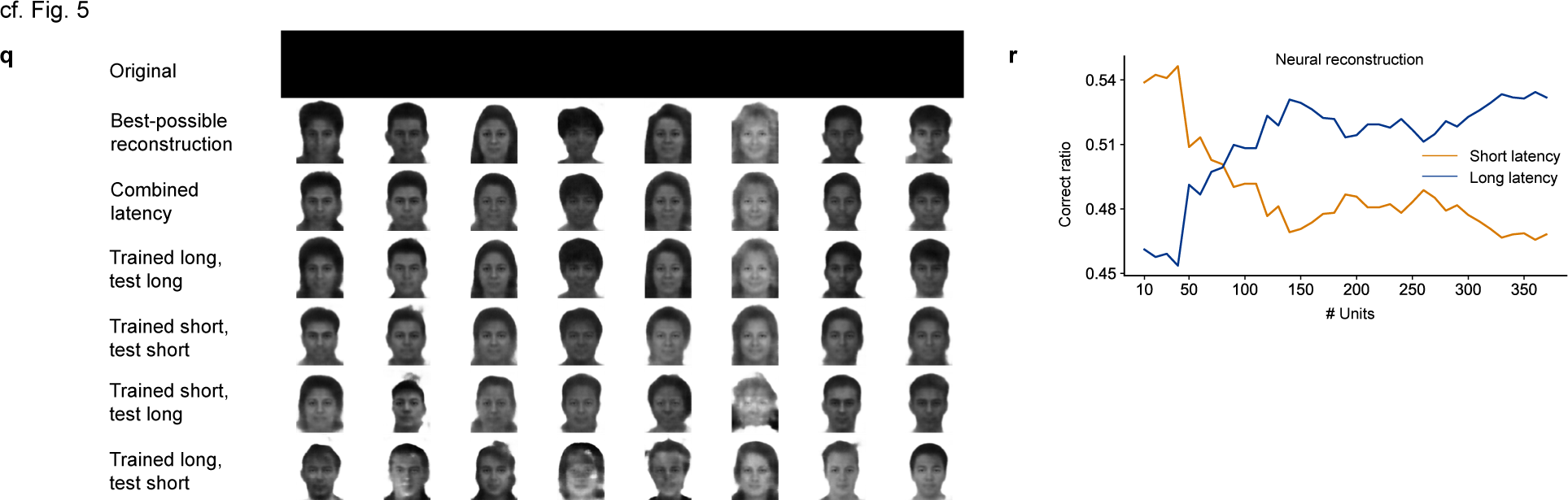
Main results computed for monkey J. [Note: all human faces in this figure were synthesized using a variational autoencoder^58^; they are not photographs. The faces in the first row of panel q have been redacted. Thus all images in this figure are in accordance with biorxiv policy on displaying human faces.] **a.** Distribution of object (left, green) and face (right, purple) axes, projected onto the top two dimensions of the 60-d object space (N = 131 cells, only cells with R^2^ > 0 on test set for object and face axes are included). **b.** Scatter plots of object vs. face axis weights for PC1 (left) and PC2 (right). **c.** Cross prediction accuracies, same conventions as Fig. 2c. **d.** Histogram of correlations between the face and object axes for each cell in the 60-d object space for real (dark gray) and AlexNet (light gray) units, normalized by the correlation between axes from two independent stimulus subsets. **e.** From left to right: matrices of mean cosine similarities across the population between (object, object), (face, face), and (face, object) axes for different pairs of latencies (N = 131 cells). **f.** Correlation between face and object axes as a function of time (diagonal values in the rightmost similarity matrix in (e)). **g.** Mean response time course to each face and object stimulus, averaged across cells and trials. **h.** Cosine similarity between the overall object axis for each cell (computed using a time window of 50-220 ms) and its time-varying face (top) and object (bottom) axes, sorted from top to bottom according to face selectivity d’ (left). **i.** Response sparsity as a function of time for faces and objects. **j.** Time-resolved scatter plots of face and object stimuli projected onto PC1 and PC2 of object space, color coded by the mean response magnitude across the population (N = 131 cells) to each stimulus. **k.** Matrix of face axis weights in AlexNet face space for each cell, computed using a short (80- 100 ms, top), long-early (120-140 ms, middle), and long-late (160-180 ms, bottom) latency window; same conventions as Fig. 4a. **l.** Matrix of object axis weights; conventions same as (k). **m.** Purple (green) distribution shows correlation coefficients across units between face (object) axis weights at short and long-early latency. Conventions as in Fig. 4c. Face axis weights were negatively corelated (one sample t-test, t(130) = −6.03, p < 1.7*10^-8^), object axis weights were positively correlated (one sample t-test, t(130) = 12.75, p < 1.3*10^-24^). **n.** Same as (m), using higher face space dimensions 6-60 to compute correlation for each cell. Face axis weights were uncorrelated (one sample t-test, t(130) = 0.52, p = 0.60), object axis weights were positively correlated (one sample t-test, t(130) = 6.96, p < 1.5*10^-10^). **o.** Same as (m), using axis weights at long-early and long-late latencies (120-140 ms and 160- 180 ms) and all 60 dimensions. Face and object axis weights were both positively correlated (face: one sample t-test, t(130) = 21.83, p < 2.6*10^-45^, object: one sample t-test, t(130) = 22.39, p < 2*10^-46^). **p.** Explained variance in z-scored population responses to faces at short (80-100 ms, orange) and long (120-140 ms, blue) latencies, as a function of number of PCs. More dimensions were required to explain 90% of response variance at long compared to short latency (97 dim for long, 83 dim for short). **q.** Reconstructions of faces using responses from different time windows. Same conventions as Fig. 5b. **r.** Face identification performance on faces reconstructed from neural activity in different time windows. Same conventions as Fig. 5d.

**Extended Data Fig. 3:**
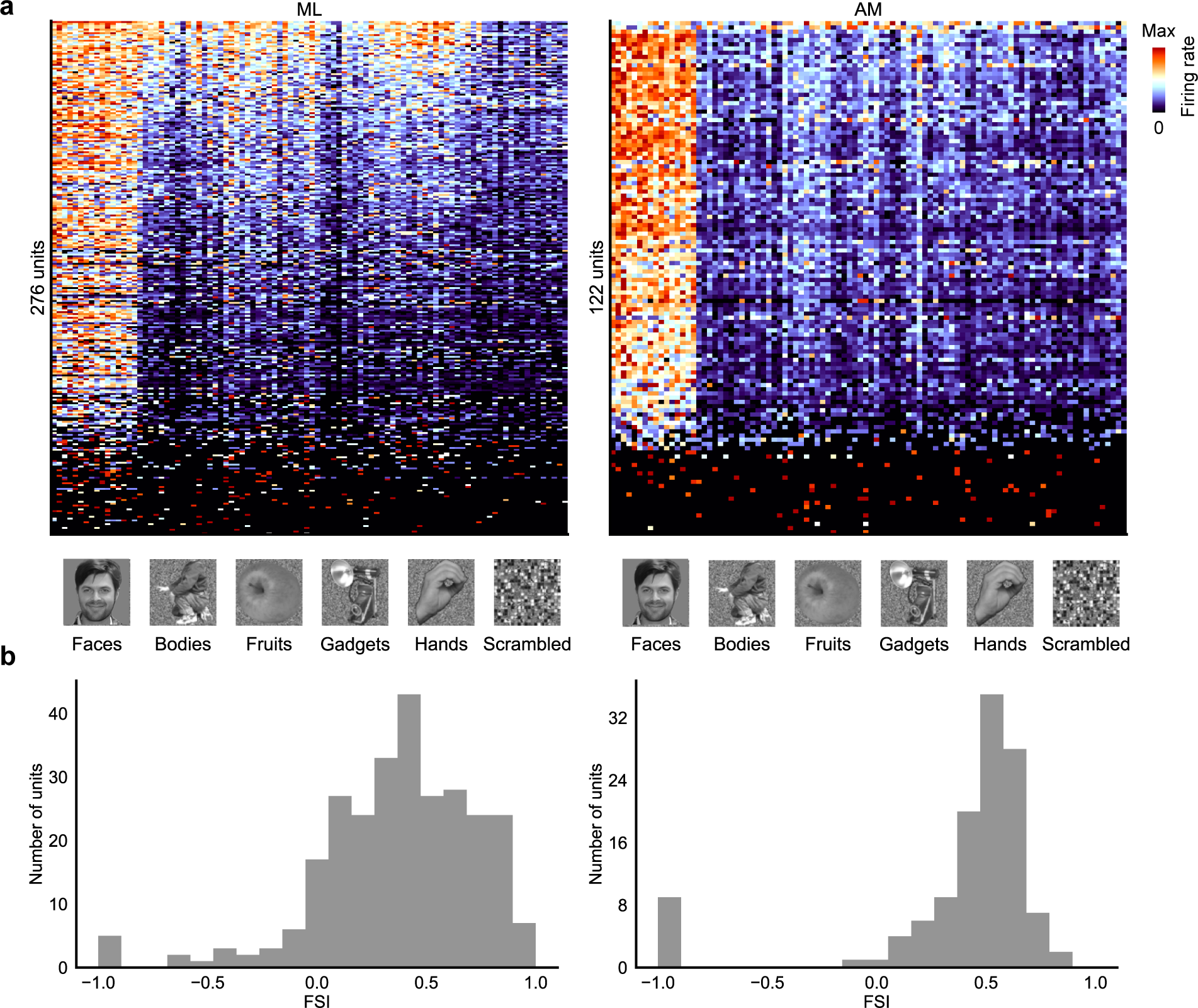
Response profiles to a screening stimulus and distributions of face selectivity indices (FSI) in ML and AM. [Note: all human faces in this figure were synthesized using a variational autoencoder^58^; they are not photographs. The faces in the first row of panel r have been redacted. Thus all images in this figure are in accordance with biorxiv policy on displaying human faces.] **a.** Mean responses of cells in ML (left) and AM (right) to 96 screening stimuli (16 images from each of the categories shown), computed using a time window of 50 – 220 ms and normalized to the range [0 1] for each cell. Only cells that were visually responsive to this stimulus set were included. **b.** Distributions of FSI (see Methods) for cells in ML (left) and AM (right). Data from monkey A.

**Extended Data Fig. 4:**
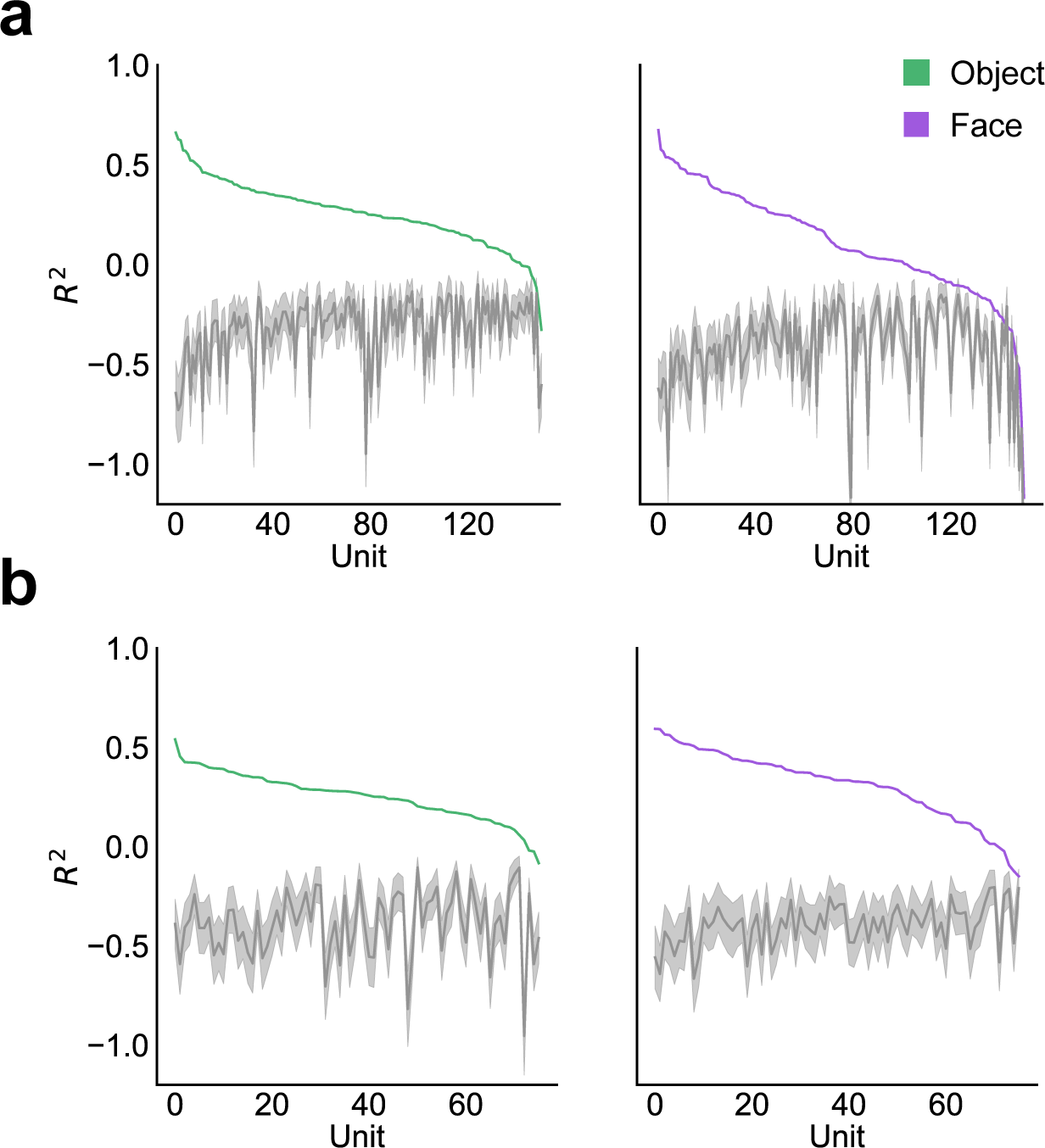
Quantification of axis tuning. **a.** Left: Explained variance of object responses for each ML cell using the object axis (green) together with explained variance for stimulus-shuffled data (gray). Right: Explained variance of face responses for each cell using the face axis (purple) together with explained variance for stimulus-shuffled data (gray). Across the population, 147/151 (151/151) units had significantly higher face (object) axis R^2^ than a random shuffle. **b.** Same as (a) for AM; 58/76 (73/76) units had significantly higher face (object) axis R^2^ than a random shuffle.

**Extended Data Fig. 5:**
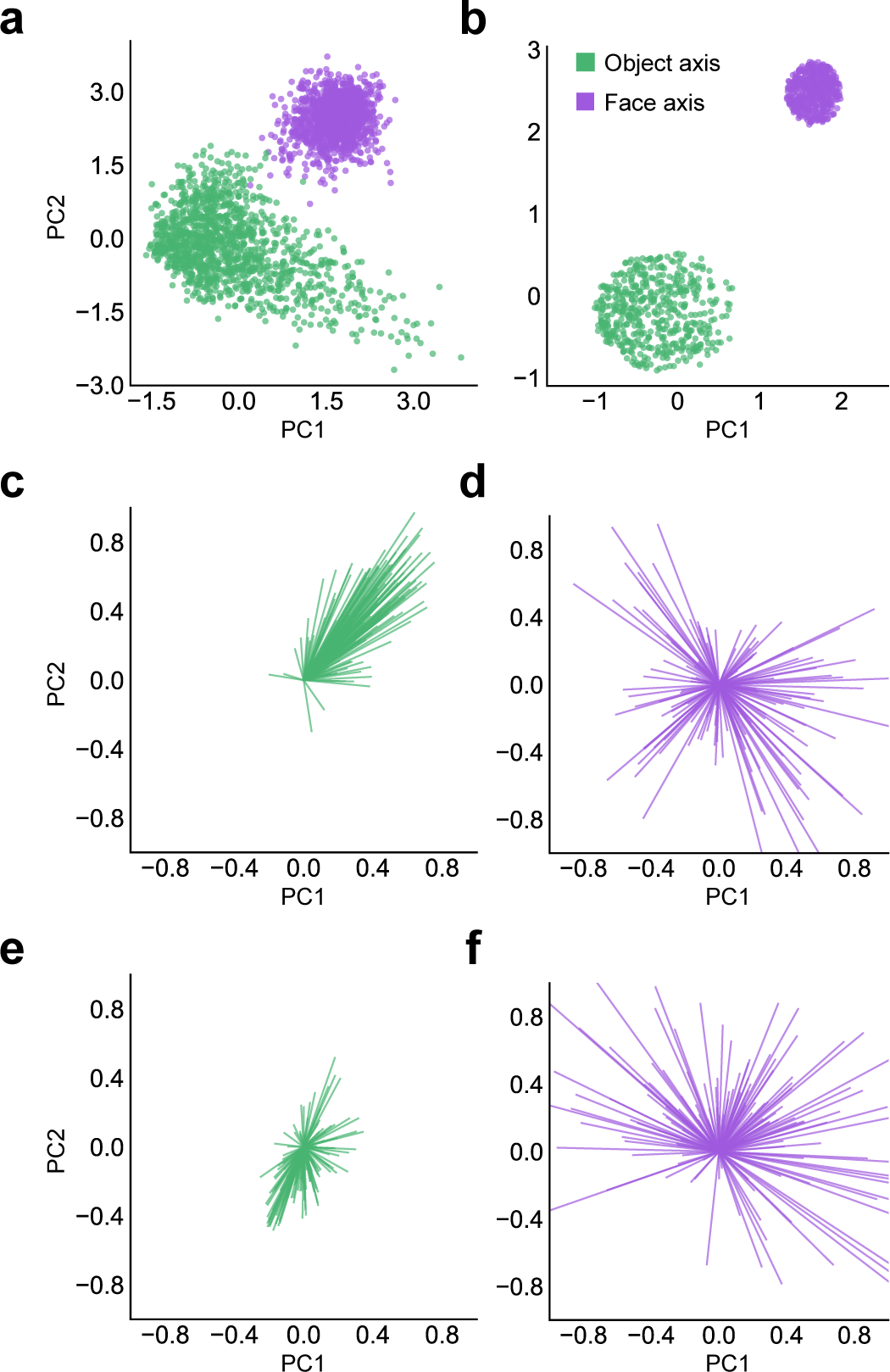
Controlling for stimulus distribution. **a.** Distribution of full set of 1425 face (purple) and 1292 object (green) stimuli projected onto the top two dimensions of the 60-d feature space. **b.** Distribution of a subset of face (purple) and object (green) stimuli chosen to make the face and object stimulus distributions maximally Gaussian. **c.** Distribution of object axes computed using responses to the subset of object images selected in (b), projected onto the top two dimensions of the 60-d feature space. **d.** Same as (c), for face axes. **e.** Same as (c), computed for stimulus-shuffled data. **f.** Same as (d), computed for stimulus-shuffled data.

**Extended Data Fig. 6:**
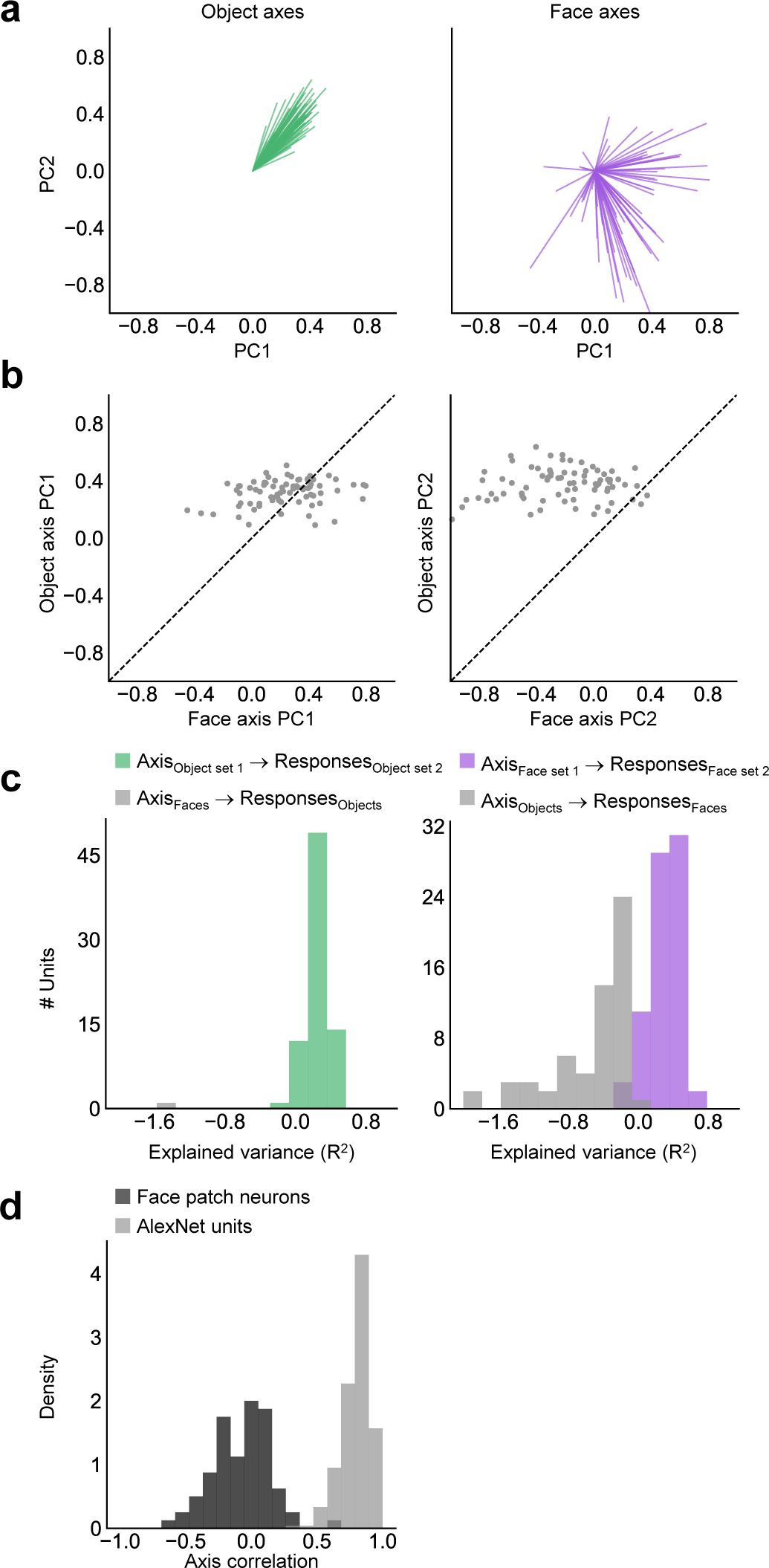
Data from face patch AM, related to Fig. 2. **a.** Distribution of object (left, green) and face (right, purple) axes, projected onto the top two dimensions of the 60-d object space (N = 76 cells, only cells with R^2^ > 0 on test set for object and face axes are included). **b.** Scatter plots of object vs. face axis weights for PC1 (left) and PC2 (right). **c.** Cross prediction accuracies, same conventions as Fig. 2c. **d.** Histogram of correlations between the face and object axes for each cell in the 60-d object space for real (dark gray) and AlexNet (light gray) units, same conventions as Extended Data Fig. 2d.

**Extended Data Fig. 7:**
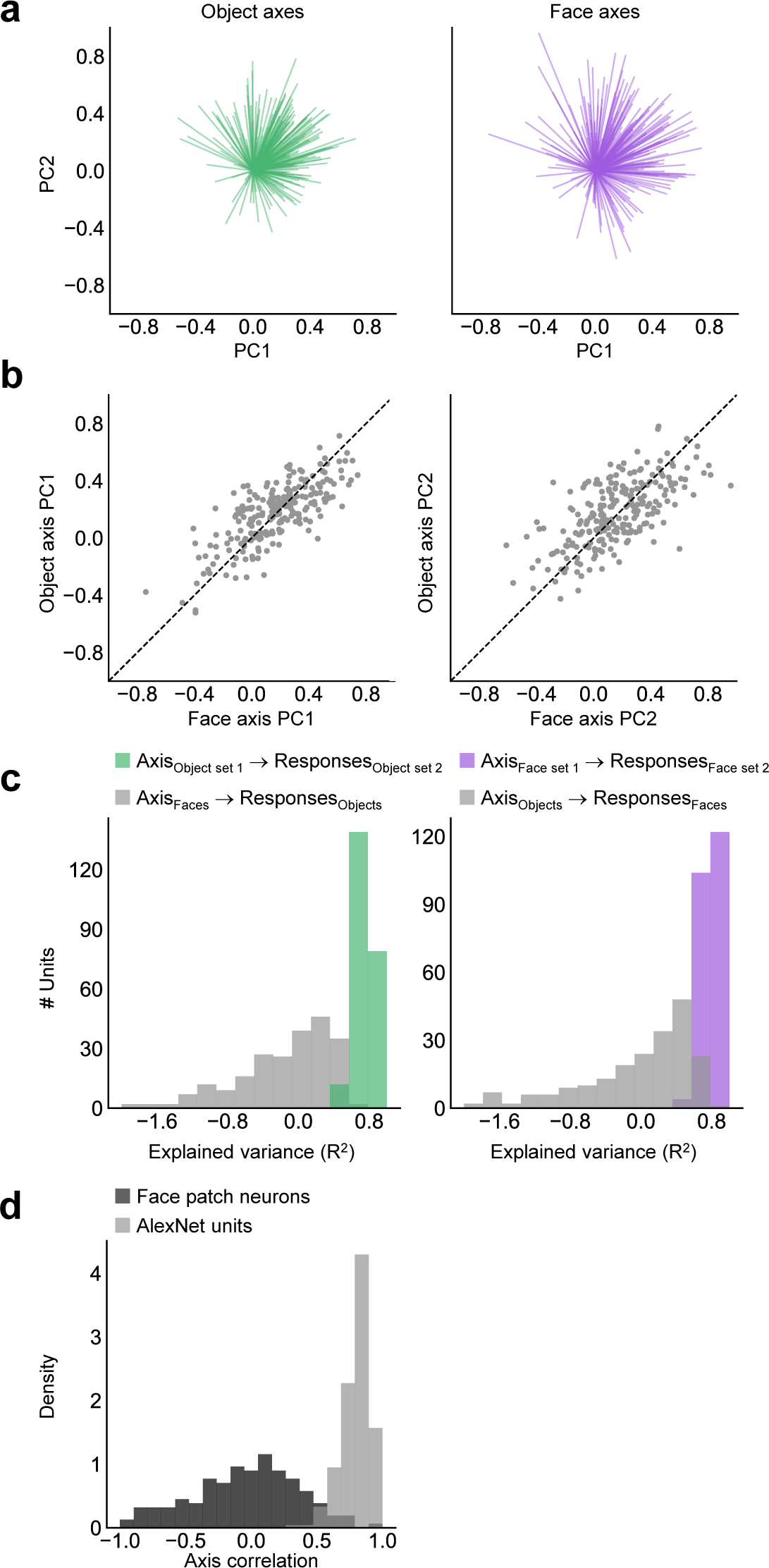
AlexNet units show correlated face and object axes. **a-c.** Same as Fig. 2a-c, computed using the 230 most-face selective units from AlexNet layer fc6 (FSI ranging from 0.2 to 0.46). **d.** Histogram of correlations between the face and object axes for each cell in the 60-d space for real (dark gray) and AlexNet (light gray) units, same conventions as Extended Data Fig. 2d.

**Extended Data Fig. 8:**
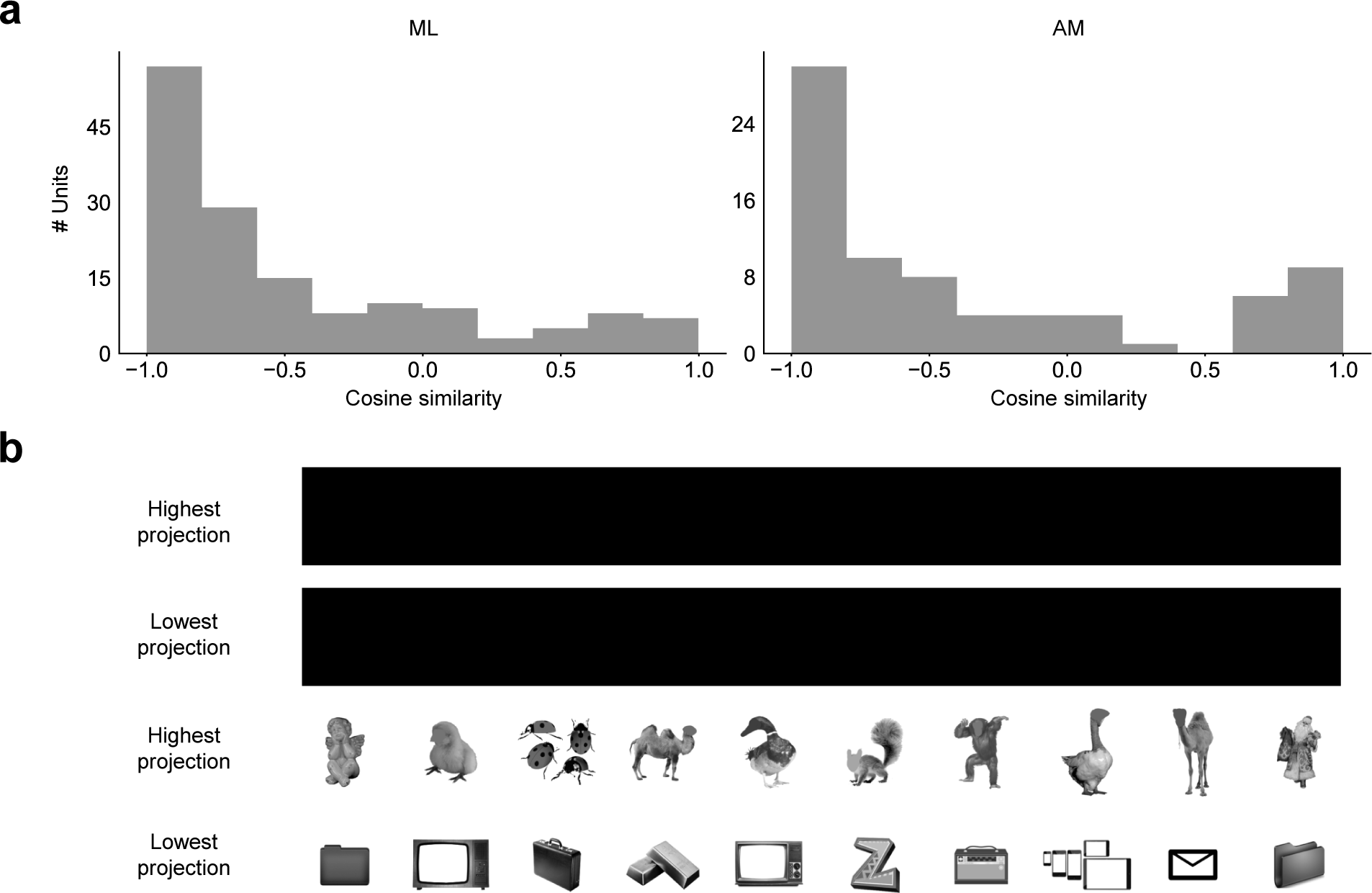
Quantification of face axis reversal. [Note: The human faces in this figure were redacted, in accordance with biorxiv policy on displaying human faces.] **a.** Distribution of cosine similarities between the first two dimensions of the face axis computed over 140-160 ms (ML) or 160-180 ms (AM) and the overall object axis computed over 50-220 ms, for ML (left) and AM (right). Cells with cosine similarity less than −0.5 (corresponding to an angle of at least 120 degrees) between the two axes were counted as reversing. **b.** Face and object images projecting maximally onto the extremes of the first two components of the mean object axis (computed in a time window of 50-220 ms and averaged across the population).

**Extended Data Fig. 9:**
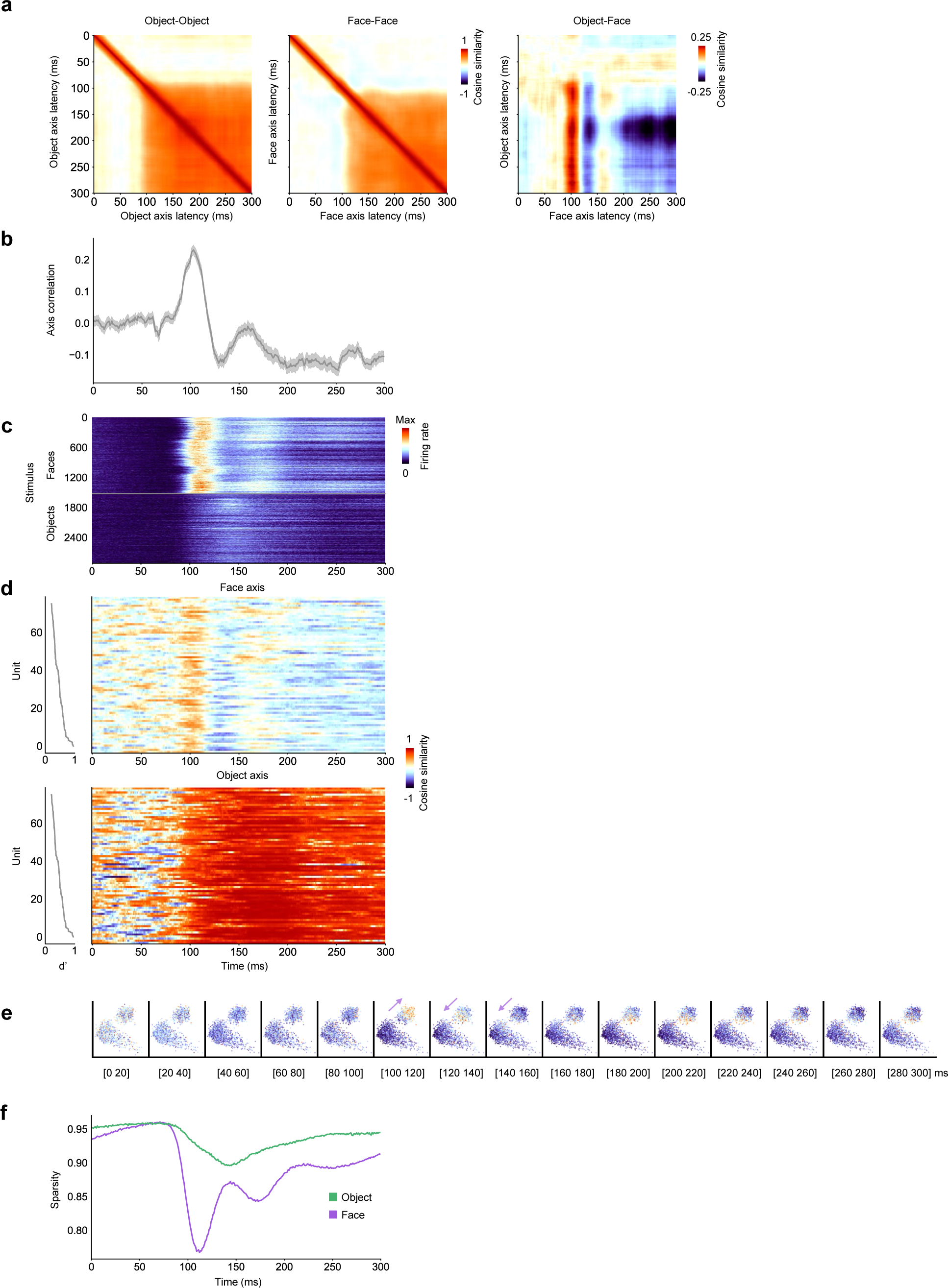
Data from face patch AM, related to Fig. 3. **a.** From left to right: matrices of mean similarities across the population between (object, object), (face, face), and (face, object) axes for different pairs of latencies (N = 76 cells). **b.** Correlation between face and object axes as a function of time (diagonal values in the rightmost similarity matrix in (a)). **c.** Mean response time course to each face and object stimulus, averaged across cells and trials. **d.** Cosine similarity between the overall object axis for each cell (computed using a time window of 50 – 220 ms) and its time-varying face (top) and object (bottom) axes, sorted from top to bottom according to d’ (left). **e.** Time-resolved scatter plots of face and object stimuli projected onto PC1 and PC2 of object space, color coded by the mean response magnitude across the population (N = 76 cells) to each stimulus. **f.** Response sparsity as a function of time for faces and objects.

**Extended Data Fig. 10:**
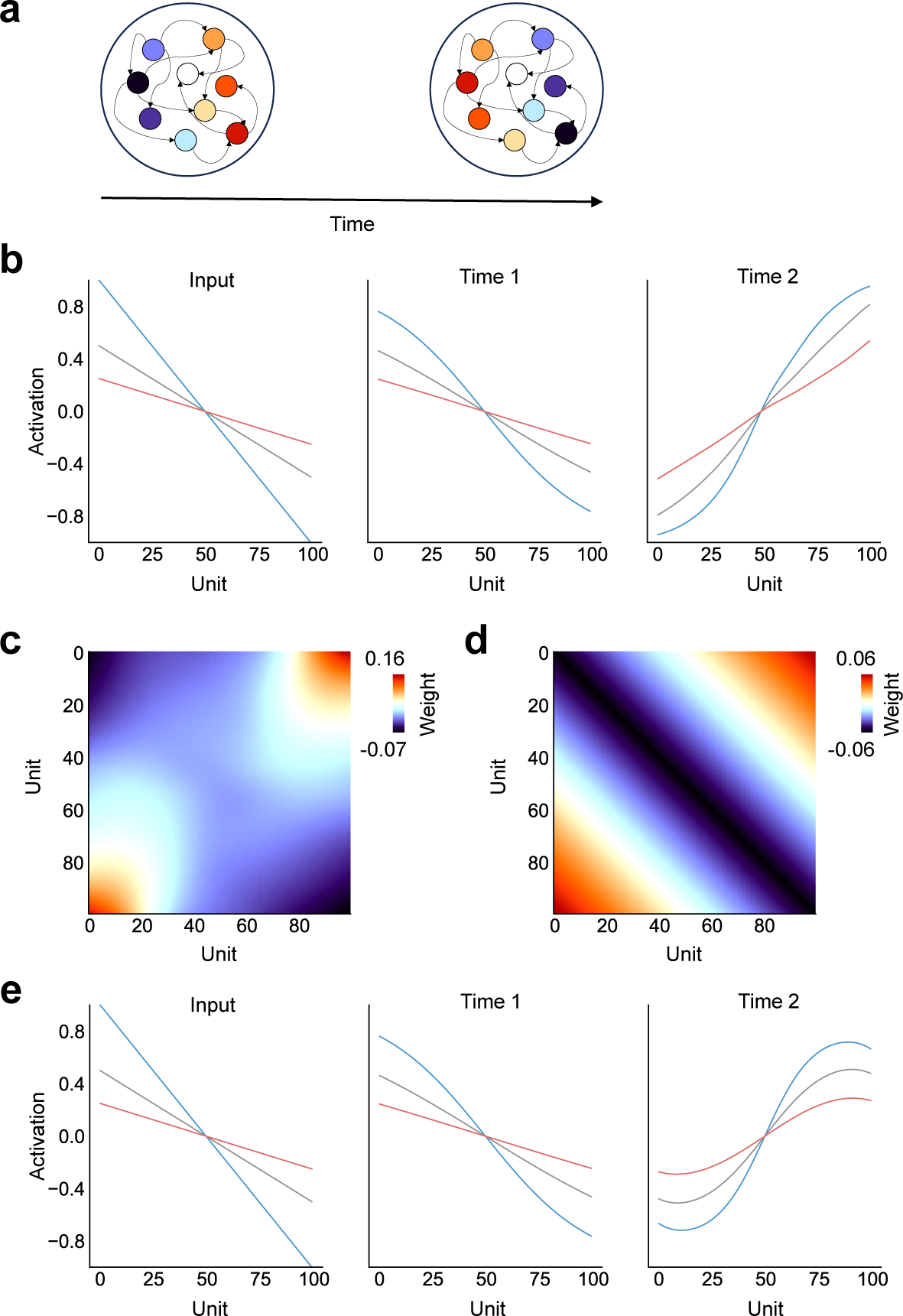
An RNN model of face axis reversal. **a.** The architecture of a simple RNN trained to reverse a gradient in its input representation (see Methods). **b.** Activity of each RNN unit over time for several different input gradients (blue, gray, red), demonstrating stable gradient reversal (cf. Fig. 3e, f, Extended Data Fig. 8). **c.** Learned RNN weight matrix. The matrix has surprisingly simple structure, with local inhibition and long-range excitation. **d.** Weight matrix of a second RNN explicitly incorporating only local inhibition and long-range excitation. **e.** Activity of each RNN unit over time for several different input gradients, for an RNN with weights as in (d).

**Extended Data Fig. 11:**
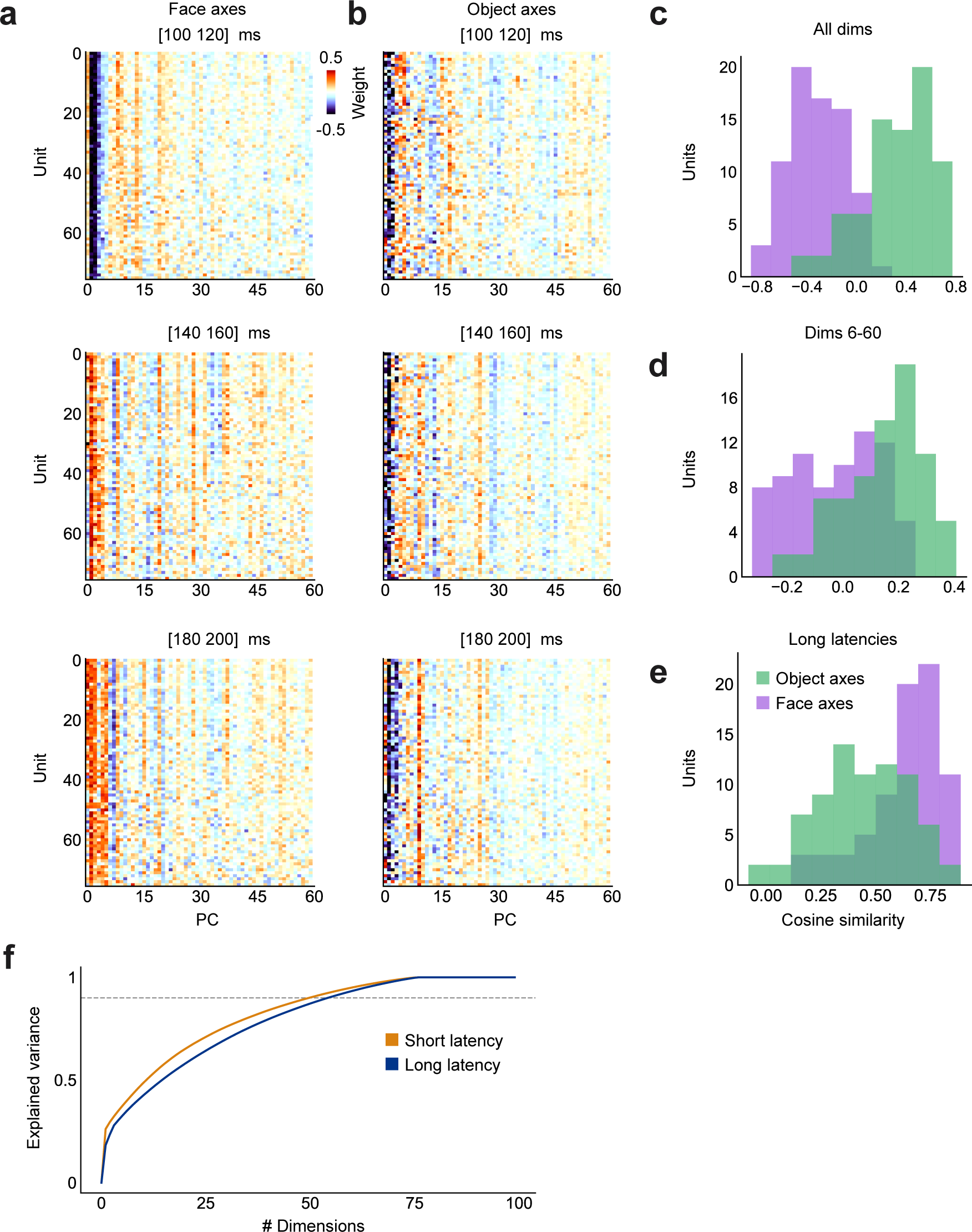
Data from face patch AM, related to Fig. 4. **a.** Matrix of face axis weights in AlexNet face space for each cell, computed using a short (80- 100 ms, top), long-early (120-140 ms, middle), and long-late (160-180 ms, bottom) latency window; same conventions as Fig. 4a. **b.** Matrix of object axis weights; conventions same as (a). **c.** Purple (green) distribution shows correlation coefficients across units between face (object) axis weights at short and long-early latency. Conventions as in Fig. 4c. Face axis weights were negatively correlated (one sample t-test, t(75) = −12.09, p < 2.7*10^-19^), object axis weights were positively correlated (one sample t-test, t(75) = 10.22, p < 7.4*10^-16^). **d.** Same as (c), using higher face space dimensions 6-60 to compute correlation for each cell. Face axis weights were not significantly correlated (one sample t-test, t(75) = −1.79, p = 0.07), object axis weights were positively correlated (one sample t-test, t(75) = 8.82, p < 3.3*10^-13^). **e.** Same as (c), using axis weights at long-early and long-late latencies (120-140 ms and 160-180 ms) and all 60 dimensions. Face and object axis weights were both positively correlated (face: one sample t-test, t(75) = 32.21, p < 1.2*10^-45^, object: one sample t-test, t(75) = 19.13, p < 1.5*10^-30^). **f.** Explained variance in population responses at short (100-120 ms, orange) and long (140-160 ms, blue) latencies, as a function of number of PCs. More dimensions were required to explain 90% of response variance at long compared to short latency (55 dim for long, 50 dim for short).

**Extended Data Fig. 12:**
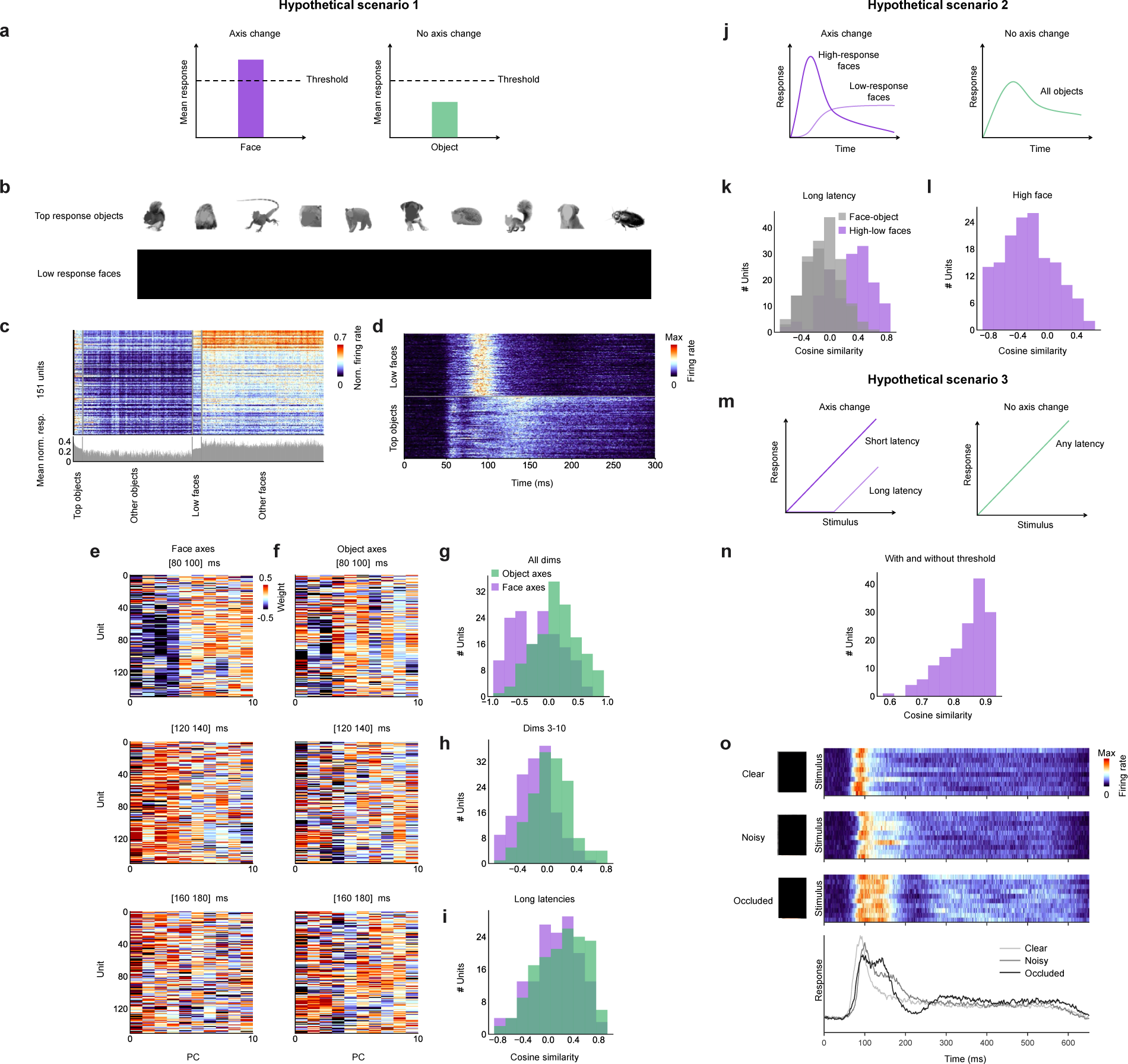
Testing three cell-intrinsic mechanisms for axis change. [Note: The human faces in this figure were redacted, in accordance with biorxiv policy on displaying human faces.] **a.** Hypothetical scenario 1: a high mean response magnitude triggers axis change. **b.** 10 most effective object and 10 least effective face stimuli. **c.** Top: Mean responses averaged over 50-220 ms to 100 most effective non-face object stimuli, other objects, 100 least effective face stimuli, and other faces. Bottom: bar graph of response to each stimulus averaged across cells (N = 151 cells). **d.** Response time courses to the 100 least effective faces (top) and 100 most effective non-face objects (bottom), averaged across cells. **e.** Matrix of face axis weights in AlexNet face space for each cell, same conventions as in Fig. 4a. Here, we only fit axes to the top 10 face space dimensions due to a small sample size (i.e., fewer images to fit the axes). **f.** Matrix of object axis weights in AlexNet face space for each cell, same conventions as in (e). **g.** Purple (green) distribution shows cosine similarities across units between face (object) axis weights at short and long-early latency; conventions as in Fig. 4c. All dimensions (1-10) were used to compute cosine similarities shown here. Face axis weights were negatively corelated (one sample t-test, t(150) = −6.60, p < 6.8*10^-10^), object axis weights were positively correlated (one sample t-test, t(150) = 4.85, p < 3.2*10-6). **h.** Same as (g), using higher face space dimensions 3-10 to compute cosine similarities for each cell. Face axis weights were negatively correlated (one sample t-test, t(150) = −7.31, p < 1.5*10^-11^), object axis weights were positively correlated (one sample t-test, t(150) = 2.55, p < 0.012). **i.** Same as (g), showing cosine similarities across units between face (object) axis weights at long-early and long-late latencies (120-140 ms and 160-180 ms), using all 10 dimensions. Face and object axis weights were both positively correlated (face: one sample t-test, t(150) = 4.13, p < 6*10^-5^, object: one sample t-test, t(150) = 6.60, p < 6.6*10^-10^). **j.** Hypothetical scenario 2: delayed responses to weaker stimuli leads to axis change. **k.** Purple distribution shows cosine similarities across units between face axis weights at long (120-140 ms) latency computed using 50% most and 50% least effective faces for each cell (see main text for details); the two sets of face axes were positively correlated (one sample t-test, t(150) = 10.58, p < 6.9*10^-20^). Gray distributions: cosine similarities between the two sets of face axes and the object axes at the same latency; the two sets of face axes were both negatively correlated with object axes (low face-object: one sample t-test, t(150) = −5.59, p < 1.1*10^-7^; high face-object: one sample t-test, t(150) = −4.52, p < 1.3*10^-5^). **l.** Distribution of cosine similarities across units between face axis weights at short (80-100 ms) and long (120-140 ms) latency computed using 50% most effective faces for each cell; axes were negatively correlated (one sample t-test, t(150) = −8.69, p < 5.6*10^-15^). **m.** Hypothetical scenario 3: cell-intrinsic adaptation leads to axis change. **n.** Distribution of cosine similarities across units between face axis weights of hypothetical units tuned to a single axis computed with and without application of a raised threshold (set to one standard deviation above mean response); axes were significantly correlated (one sample t-test, t(150)=146.67, p < 7*10^-164^). **o.** Response time courses to clear (Row 1), noisy (Row 2), and occluded (Row 3) faces (10 identities each), averaged across cells. The stimulus-averaged response time course is shown in Row 4.

**Extended Data Fig. 13:**
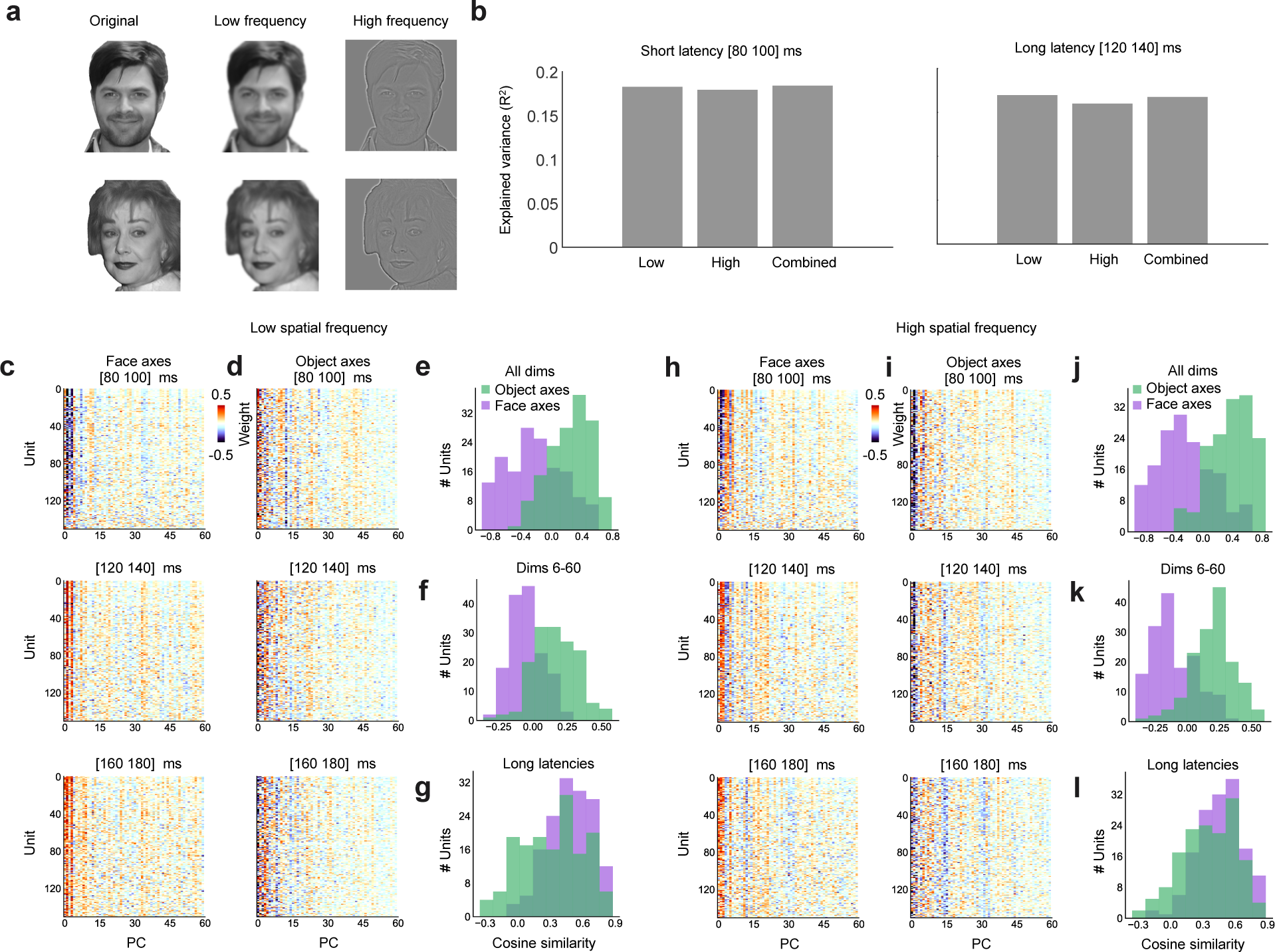
Comparing contributions of low and high spatial frequency components to axis change. [Note: The human faces in this figure were synthesized using a diffusion model^57^ and are not photographs, in accordance with biorxiv policy on displaying human faces. These stimuli were not actually shown to the animal.] **a.** Two example faces (left) filtered to show low (middle) and high (right) spatial frequency components (see Methods). **b.** Explained variance for short (left, 80-100 ms) and long (right, 120-140 ms) latency responses, using low spatial frequency features, high spatial frequency features, or a combined set of features. The number of features used to compute explained variance was the same in all three conditions (low: 120, high: 120, combined: 60+60). **c-g.** Axis weights and axis weight correlations computed using low spatial frequency features, same conventions as Fig. 4a-e. Face axis weights showed negative correlation between the two time windows (all dimensions: one sample t-test, t(150) = −7.47, p < 6.3*10^-12^; higher dimensions: one sample t-test, t(150) = −4.56, p < 1.1*10^-5^). Object axis weights were positively correlated between the two time windows (all dimensions: one sample t-test, t(150) = 16.07, p < 2.1*10^-34^; higher dimensions: one sample t-test, t(150) = 12.69, p < 1.6*10^-25^). Both face and object axis weights were positively correlated between the two long latency windows (face: one sample t-test, t(150) = 29.28, p < 6.7*10^-64^; object: one sample t-test, t(150) = 13.70, p < 3.2*10^-28^). **h-l.** Axis weights and axis weight correlations computed using high spatial frequency features, same conventions as Fig. 4a-e. Face axis weights showed negative correlation between the two time windows (all dimensions: one sample t-test, t(150) = --8.05, p < 2.4*10^-13^; higher dimensions: one sample t-test, t(150) = −7.59, p < 3.2*10^-12^). Object axis weights were positively correlated between the two time windows (all dimensions: one sample t-test, t(150) = 16.07, p < 2.1*10^-34^; higher dimensions: one sample t-test, t(150) = 14.84, p < 3.1*10^-31^). Both face and object axis weights were positively correlated between the two long latency windows (face: one sample t-test, t(150) = 26.80, p < 4.6*10^-59^; object: one sample t-test, t(150) = 17.50, p < 4.6*10^-38^).

**Extended Data Fig. 14:**
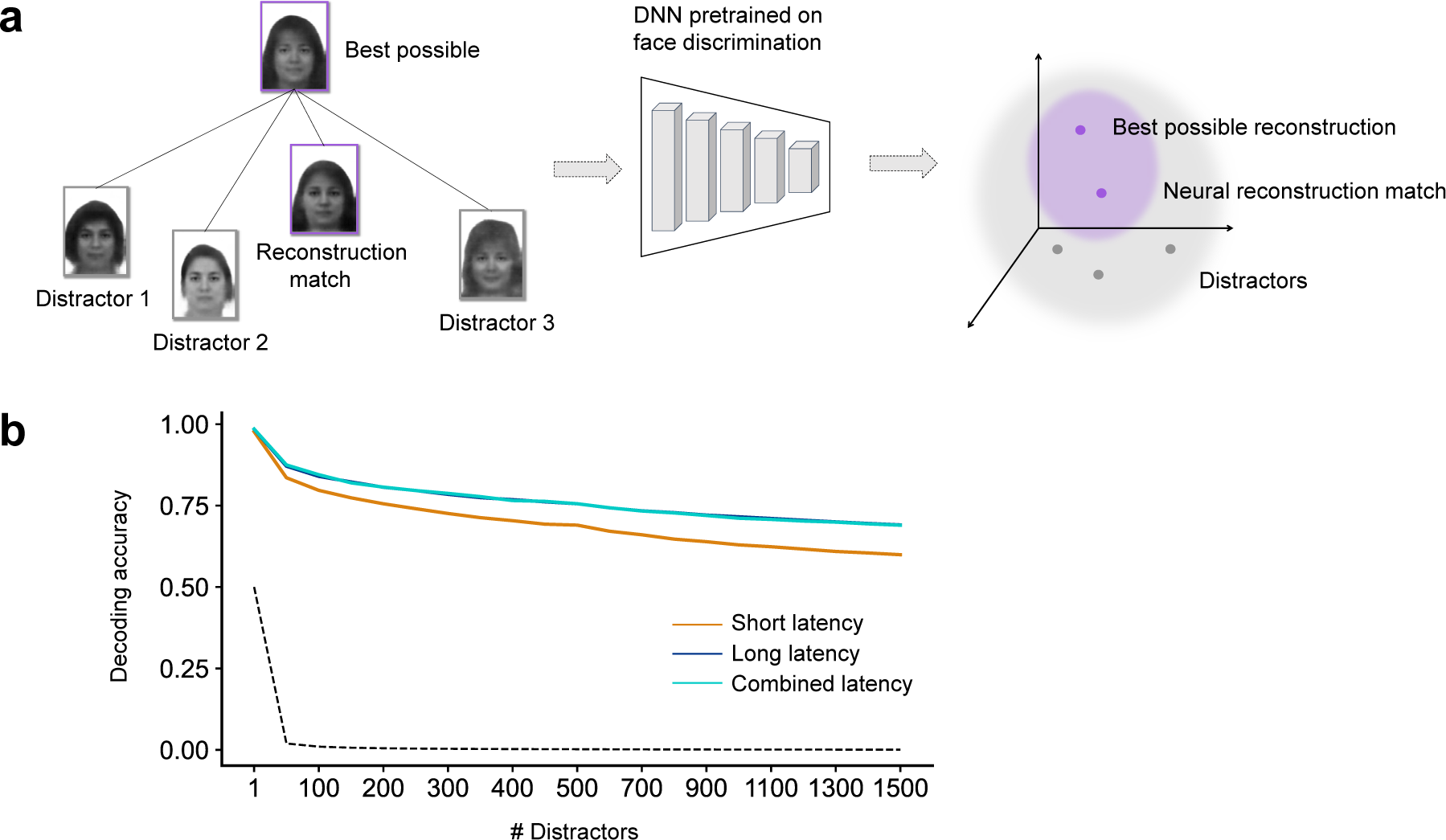
Quantifying accuracy for identifying reconstructed faces among distractors. [Note: all human faces in this figure were synthesized using a variational autoencoder^58^ and do not depict any real human, in accordance with biorxiv policy on displaying human faces.] **a.** Schematic showing the use of a DNN trained on face discrimination to test identification performance for reconstructed faces. A face was considered identified correctly if it was closer in the DNN’s face space to the optimally reconstructed face than any distractor. **b.** Face identification performance on faces reconstructed from neural activity in different time windows as a function of # distractors, following scheme in (a). The shaded area denotes standard error of the mean. Dotted line shows chance performance.

## References

1. Hubel, D. H. & Wiesel, T. N. Receptive fields, binocular interaction and functional architecture in the cat’s visual cortex. J. Physiol. 160, 106–154 (1962).

2. Yamins, D. L. K. et al. Performance-optimized hierarchical models predict neural responses in higher visual cortex. Proceedings of the National Academy of Sciences 111, 8619–8624 (2014).

3. Chang, L. & Tsao, D. Y. The Code for Facial Identity in the Primate Brain. Cell 169, 1013–1028.e14 (2017).

4. Ponce, C. R. et al. Evolving Images for Visual Neurons Using a Deep Generative Network Reveals Coding Principles and Neuronal Preferences. Cell 177, 999–1009.e10 (2019).

5. Kar, K., Kubilius, J., Schmidt, K., Issa, E. B. & DiCarlo, J. J. Evidence that recurrent circuits are critical to the ventral stream’s execution of core object recognition behavior. Nat. Neurosci. 22, 974–983 (2019).

6. Bao, P., She, L., McGill, M. & Tsao, D. Y. A map of object space in primate inferotemporal cortex. Nature 583, 103–108 (2020).

7. Gross, C. G., Rocha-Miranda, C. E. & Bender, D. B. Visual properties of neurons in inferotemporal cortex of the Macaque. J. Neurophysiol. 35, 96–111 (1972).

8. Mishkin, M., Ungerleider, L. G. & Macko, K. A. Object vision and spatial vision: two cortical pathways. Trends Neurosci. 6, 414–417 (1983).

9. Tanaka, K. Inferotemporal cortex and object vision. Annu. Rev. Neurosci. 19, 109–139 (1996).

10. Verhoef, B. E., Vogels, R. & Janssen, P. Inferotemporal cortex subserves three-dimensional structure categorization. Neuron 73, 171–182 (2012).

11. Lafer-Sousa, R. & Conway, B. R. Parallel, multi-stage processing of colors, faces and shapes in macaque inferior temporal cortex. Nat. Neurosci. 16, 1870–1878 (12 2013).

12. Vaziri, S., Carlson, E. T., Wang, Z. & Connor, C. E. A Channel for 3D Environmental Shape in Anterior Inferotemporal Cortex. Neuron 84, 55–62 (2014).

13. Popivanov, I. D., Jastorff, J., Vanduffel, W. & Vogels, R. Heterogeneous single-unit selectivity in an fMRI-defined body-selective patch. J. Neurosci. 34, 95–111 (2014).

14. Tsao, D. Y., Freiwald, W. A., Knutsen, T. A., Mandeville, J. B. & Tootell, R. B. H. Faces and objects in macaque cerebral cortex. Nat. Neurosci. 6, 989–995 (2003).

15. Tsao, D. Y., Freiwald, W. A., Tootell, R. B. H. & Livingstone, M. S. A cortical region consisting entirely of face-selective cells. Science 311, 670–674 (2006).

16. Hesse, J. K. & Tsao, D. Y. The macaque face patch system: a turtle’s underbelly for the brain. Nat. Rev. Neurosci. 21, 695–716 (2020).

17. Kanwisher, N., McDermott, J. & Chun, M. M. The fusiform face area: a module in human extrastriate cortex specialized for face perception. J. Neurosci. 17, 4302–4311 (1997).

18. Hirschfeld, L. A. & Gelman, S. A. Mapping the Mind: Domain Specificity in Cognition and Culture. (Cambridge University Press, 1994).

19. Yovel, G. & Kanwisher, N. Face perception: domain specific, not process specific. Neuron 44, 889–898 (2004).

20. Haxby, J. V. et al. Distributed and overlapping representations of faces and objects in ventral temporal cortex. Science 293, 2425–2430 (2001).

21. Gauthier, I., Tarr, M. J., Anderson, A. W., Skudlarski, P. & Gore, J. C. Activation of the middle fusiform “face area” increases with expertise in recognizing novel objects. Nat. Neurosci. 2, 568–573 (1999).

22. Vinken, K., Prince, J. S., Konkle, T. & Livingstone, M. S. The neural code for “face cells” is not face-specific. Sci Adv 9, eadg1736 (2023).

23. Tsao, D. Y. & Livingstone, M. S. Mechanisms of face perception. Annu. Rev. Neurosci. 31, 411–437 (2008).

24. Thompson, P. Margaret Thatcher: a new illusion. Perception 9, 483–484 (1980).

25. Tanaka, J. W. & Farah, M. J. Parts and wholes in face recognition. Q. J. Exp. Psychol. A 46, 225–245 (1993).

26. Bodamer, J. Prosop’s agnosia; the agnosia of cognition. Arch. Psychiatr. Nervenkr. Z. Gesamte Neurol. Psychiatr. 118, 6–53 (1947).

27. Damasio, A. R., Damasio, H. & Van Hoesen, G. W. Prosopagnosia: anatomic basis and behavioral mechanisms. Neurology 32, 331–341 (1982).

28. Freiwald, W. A. & Tsao, D. Y. Functional compartmentalization and viewpoint generalization within the macaque face-processing system. Science 330, 845–851 (2010).

29. Issa, E. B. & DiCarlo, J. J. Precedence of the eye region in neural processing of faces. J. Neurosci. 32, 16666–16682 (2012).

30. Moeller, S., Crapse, T., Chang, L. & Tsao, D. Y. The effect of face patch microstimulation on perception of faces and objects. Nat. Neurosci. 20, 743–752 (2017).

31. Ohayon, S., Freiwald, W. A. & Tsao, D. Y. What makes a cell face selective? The importance of contrast. Neuron 74, 567–581 (2012).

32. Konkle, T. & Alvarez, G. A. A self-supervised domain-general learning framework for human ventral stream representation. Nat. Commun. 13, 491 (2022).

33. Krizhevsky, A., Sutskever, I. & Hinton, G. E. Imagenet classification with deep convolutional neural networks. Adv. Neural Inf. Process. Syst. 25, 1097–1105 (2012).

34. Chang, L., Egger, B., Vetter, T. & Tsao, D. Y. Explaining face representation in the primate brain using different computational models. Curr. Biol. 31, 2785–2795.e4 (2021).

35. Cadieu, C. F. et al. Deep neural networks rival the representation of primate IT cortex for core visual object recognition. PLoS Comput. Biol. 10, e1003963 (2014).

36. Schrimpf, M. et al. Brain-Score: Which Artificial Neural Network for Object Recognition is most Brain-Like? bioRxiv preprint (2018).

37. DiCarlo, J. J., Zoccolan, D. & Rust, N. C. How does the brain solve visual object recognition? Neuron 73, 415–434 (2012).

38. Trautmann, E. M., et al. Large-scale high-density brain-wide neural recording in nonhuman primates. bioRxiv (2023) doi:10.1101/2023.02.01.526664.

39. Isaacson, J. S. & Scanziani, M. How inhibition shapes cortical activity. Neuron 72, 231–243 (2011).

40. Hartline, H. K., Wagner, H. G. & Ratliff, F. Inhibition in the eye of Limulus. J. Gen. Physiol. 39, 651–673 (1956).

41. Ben-Yishai, R., Bar-Or, R. L. & Sompolinsky, H. Theory of orientation tuning in visual cortex. Proc. Natl. Acad. Sci. U. S. A. 92, 3844–3848 (1995).

42. Bar, M. Visual objects in context. Nat. Rev. Neurosci. 5, 617–629 (2004).

43. Sripati, A. P. & Olson, C. R. Representing the forest before the trees: a global advantage effect in monkey inferotemporal cortex. J. Neurosci. 29, 7788–7796 (2009).

44. Livingstone, M. & Hubel, D. Segregation of form, color, movement, and depth: anatomy, physiology, and perception. Science 240, 740–749 (1988).

45. Liu, Z. et al. A ConvNet for the 2020s. arXiv [cs.CV] 11976–11986 (2022).

46. Parkhi, O.m., vedaldi, A. and Zisserman, A. (2015) Deep Face Recognition. Proceedings of the British Machine Vision Conference (BMVC). - references - scientific research publishing. https://www.scirp.org/(S(lz5mqp453edsnp55rrgjct55.))/reference/referencespapers.aspx?referenceid=2076487.

47. Sugase, Y., Yamane, S., Ueno, S. & Kawano, K. Global and fine information coded by single neurons in the temporal visual cortex. Nature 400, 869–873 (1999).

48. Brincat, S. L. & Connor, C. E. Dynamic shape synthesis in posterior inferotemporal cortex. Neuron 49, 17–24 (2006).

49. Tanigawa, H., Wang, Q. & Fujita, I. Organization of horizontal axons in the inferior temporal cortex and primary visual cortex of the macaque monkey. Cereb. Cortex 15, 1887–1899 (2005).

50. Grimaldi, P., Saleem, K. S. & Tsao, D. Anatomical Connections of the Functionally Defined “Face Patches” in the Macaque Monkey. Neuron 90, 1325–1342 (2016).

51. She, L., Benna, M. K., Shi, Y., Fusi, S. & Tsao, D. Y. The geometry of face memory. bioRxiv (2021) doi:10.1101/2021.03.12.435023.

52. Moscovitch, M., Winocur, G. & Behrmann, M. What Is Special about Face Recognition? Nineteen Experiments on a Person with Visual Object Agnosia and Dyslexia but Normal Face Recognition. J. Cogn. Neurosci. 9, 555–604 (1997).

53. Freiwald, W. A., Tsao, D. Y. & Livingstone, M. S. A face feature space in the macaque temporal lobe. Nat. Neurosci. 12, 1187–1196 (2009).

54. Arcaro, M. J., Ponce, C. & Livingstone, M. The neurons that mistook a hat for a face. Elife 9, (2020).

55. Bashivan, P., Kar, K. & DiCarlo, J. J. Neural population control via deep image synthesis. Science 364, eaav9436 (2019).

56. Tanaka, K. Columns for complex visual object features in the inferotemporal cortex: clustering of cells with similar but slightly different stimulus selectivities. Cereb. Cortex 13, 90–99 (2003).

57. Huang, Z., Chan, K. C. K., Jiang, Y. & Liu, Z. Collaborative Diffusion for multi-modal face generation and editing. arXiv [cs.CV] (2023).

58. Tolstikhin, I., Bousquet, O., Gelly, S. & Schoelkopf, B. Wasserstein Auto-Encoders. arXiv [stat.ML] (2017).

## Methods References

1. She, L., Benna, M. K., Shi, Y., Fusi, S. & Tsao, D. Y. The geometry of face memory. bioRxiv (2021) doi:10.1101/2021.03.12.435023.

2. Phillips, P. J., Wechsler, H., Huang, J. & Rauss, P. J. The FERET database and evaluation procedure for face-recognition algorithms. Image Vis. Comput. 16, 295–306 (1998).

3. Phillips, P. J., Moon, H., Rauss, P. J. & Rizvi, S. The FERET Evaluation Methodology for Face Recognition Algorithms. IEEE TPAMI 22, 1090–1104 (2000).

4. Solina, F., Peer, P., Batagelj, B., Juvan, S. & Kovac, J. Computer Graphics Collaboration for Model-based Imaging, Rendering, image Analysis and Graphical special Effects. in (2003).

5. Strohminger, N. et al. The MR2: A multi-racial, mega-resolution database of facial stimuli.Behav. Res. Methods 48, 1197–1204 (2016).

6. Ma, D. S., Correll, J. & Wittenbrink, B. The Chicago face database: A free stimulus set of faces and norming data. Behav. Res. Methods 47, 1122–1135 (2015).

7. Yang, S., Luo, P., Loy, C. C. & Tang, X. WIDER FACE: A face detection benchmark. in 2016 IEEE Conference on Computer Vision and Pattern Recognition (CVPR) 5525–5533 (IEEE, 2016).

8. Bao, P., She, L., McGill, M. & Tsao, D. Y. A map of object space in primate inferotemporal cortex. Nature 583, 103–108 (2020).

9. Ohayon, S., Freiwald, W. A. & Tsao, D. Y. What makes a cell face selective? The importance of contrast. Neuron 74, 567–581 (2012).

10. Reuter, M. & Fischl, B. Avoiding asymmetry-induced bias in longitudinal image processing. Neuroimage 57, 19–21 (2011).

11. Trautmann, E. M., et al. Large-scale high-density brain-wide neural recording in nonhuman primates. bioRxiv (2023) doi:10.1101/2023.02.01.526664.

12. Jun, J. J. et al. Fully integrated silicon probes for high-density recording of neural activity. Nature 551, 232–236 (2017).

13. Pachitariu, M., Sridhar, S. & Stringer, C. Solving the spike sorting problem with Kilosort. bioRxiv 2023.01.07.523036 (2023) doi:10.1101/2023.01.07.523036.

14. Ohayon, S. & Tsao, D. Y. MR-guided stereotactic navigation. Journal of Neuroscience Methods vol. 204 389–397 Preprint at 10.1016/j.jneumeth.2011.11.031 (2012).

15. Krizhevsky, A., Sutskever, I. & Hinton, G. E. Imagenet classification with deep convolutional neural networks. Adv. Neural Inf. Process. Syst. 25, 1097–1105 (2012).

16. Willmore, B. & Tolhurst, D. J. Characterizing the sparseness of neural codes. Network 12, 255–270 (2001).

17. Tolstikhin, I., Bousquet, O., Gelly, S. & Schoelkopf, B. Wasserstein Auto-Encoders. arXiv [stat.ML] (2017).

18. Kingma, D. P. & Welling, M. Auto-encoding variational bayes. arXiv preprint arXiv:1312. 6114 (2013).

19. Liu, Z. et al. A ConvNet for the 2020s. arXiv [cs.CV] 11976–11986 (2022).

